# Brown fat protects against hepatic oxidative stress by remodeling the circulating metabolome

**DOI:** 10.64898/2026.05.12.722834

**Authors:** Dandan Wang, Mark Li, Tian Lu, Mami Matsushita, Juro Sakai, Masayuki Saito, Takeshi Yoneshiro, Shingo Kajimura

## Abstract

Brown adipose tissue (BAT) regulates systemic metabolism beyond thermogenesis, yet the circulating mediators through which BAT communicates with other organs remain less explored. Here, we performed comprehensive serum metabolomics and lipidomics in BAT-ablated mice and human cohorts with varying BAT activity to delineate how BAT activity shapes the circulating metabolome. By integrating datasets across serum, tissues, extracellular fluids, and conditioned media, we assembled BAT-linked circulating molecular signatures. The analyses support a critical role for BAT in the clearance of circulating branched-chain amino acids and triglycerides. We also identified a cold-inducible metabolite, 3-hydroxystearic acid (3-OHSA), produced primarily by BAT and released into circulation. 3-OHSA serves as a circulating readout of cold-activated BAT and acts on the liver to reduce mitochondrial membrane potential and reactive oxygen species production, thereby limiting oxidative stress. This work provides a framework for identifying BAT-derived mediators and uncovers a BAT-liver axis that coordinates adaptation to metabolic stress.

## INTRODUCTION

Brown adipose tissue (BAT) is a metabolically active organ that protects against cardiometabolic diseases in humans and rodents ^1–5^. Because BAT is historically defined as a thermogenic organ through the action of a BAT-specific protein, uncoupling protein 1 (UCP1), the prevailing model holds that UCP1-mediated heat production increases whole-body energy expenditure, reduces body weight, and thereby improves glucose homeostasis and insulin sensitivity. Although many animal studies support this linear framework, several observations challenged the notion. For example, adipose-specific loss of key brown-fat regulators (e.g., the PRDM16-EHMT1 complex) led to hepatic insulin resistance, adipose fibrosis, and inflammation even at ambient temperature prior to changes in body weight ^6–8^, whereas UCP1-deficient mice do not develop obesity or insulin resistance unless housed at thermoneutrality ^9–11^. Furthermore, a BAT-selective defect in branched-chain amino acid (BCAA) catabolism resulted in elevated hepatic oxidative stress and systemic insulin resistance, despite preserved BAT thermogenesis and whole-body energy expenditure ^12^. Conversely, a recent study showed that enhanced browning of white adipose tissue (WAT) by the fat-specific deletion of RIP140 led to improved hepatic insulin sensitivity without altering body weight or energy expenditure ^13^. Importantly, large-scale human cohort studies showed that active BAT is significantly associated with lower prevalences of Type 2 diabetes, dyslipidemia, and cardiovascular disease – all of which remain significant independent of body mass index (BMI) ^14–16^. Together, these findings indicate that the metabolic benefits of BAT extend far beyond its well-defined thermogenic action and whole-body energy expenditure ^17,18^. A key question is what mediates the metabolically favorable action of BAT in controlling metabolic health.

Converging evidence suggests that BAT communicates systemically by releasing peptides, microRNAs, metabolites, and lipids, collectively termed batokines, which act in endocrine, paracrine, and/or autocrine manners ^19–21^. For example, fibroblast growth factor 21 (FGF21) is induced by cold and can be secreted by BAT to promote glucose utilization in adipose tissue, although the majority of circulating FGF21 originates from the liver ^22,23^. A cold-inducible lipid, 12,13-dihydroxy-9Z-octadecenoic acid (12,13-diHOME), is released from BAT as well as skeletal muscle, which enhances fatty-acid uptake in BAT ^24,25^. Secretome profiling of human brown adipocytes identified ependymin-related protein-1 (EPDR1) as a protein linked to thermogenesis and glucose-stimulated insulin secretion in beta cells, while it is also expressed in multiple tissues ^26,27^. These studies underscore BAT’s endocrine function to communicate with other metabolic organs via batokines. Meanwhile, the studies raise a key challenge for the field: defining circulating biomarkers that reliably reflect BAT activity, since ^18^F-fluoro-2-deoxy-D-glucose (^18^F-FDG) PET/CT imaging, the current gold standard, is costly, exposes patients to radiation, and limits large-scale application^28^.

Here, we attempted to address this gap by employing a circulation-first approach, *i.e.,* prioritizing the contribution of BAT to the circulating metabolite landscape *in vivo*, rather than starting with the search for secreted products from cultured adipocytes or isolated BAT (**Figure 1A**). In brief, we comprehensively profiled the circulating metabolome and lipidome in mice with genetic ablation of BAT to define BAT-linked metabolites and lipid species released by or cleared through BAT *in vivo*. We then asked which of these metabolites are associated with human BAT activity, assessed by ^18^F-FDG PET/CT. By integrating mouse and human datasets, along with multi-level analyses across tissues, extracellular fluids, and conditioned media, we assembled BAT-linked circulating molecular signatures and provided mechanistic insights into how BAT mediates inter-organ crosstalk beyond thermogenesis.

**Figure 1.**
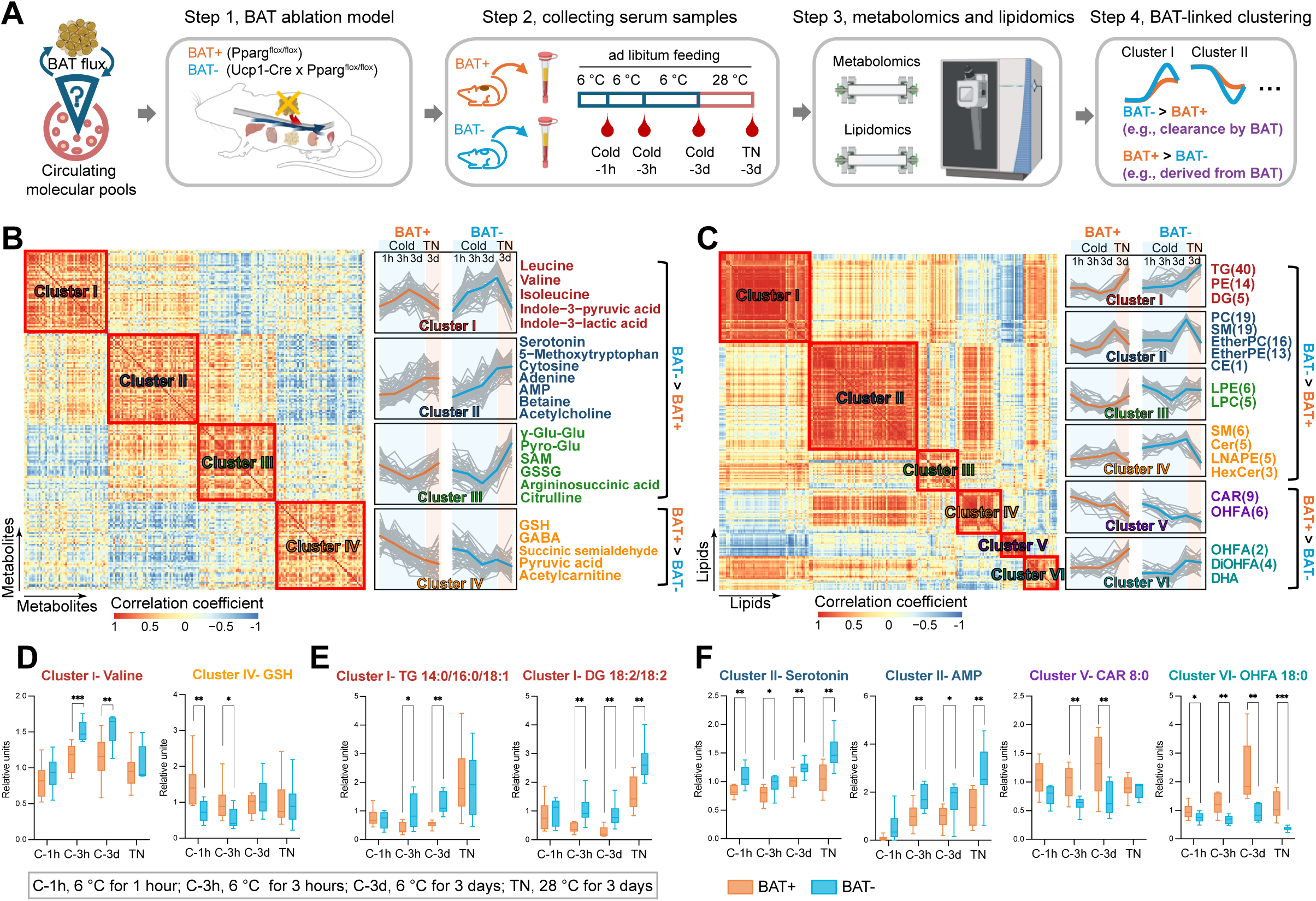
The impact of BAT on circulating metabolome and lipidome *in vivo*. **A.** Schematic overview of the strategy we used to study how BAT affects systemic metabolite and lipid profiles *in vivo. n* = 8 for BAT-ablated mice and *n* = 10 for littermate controls. **B-C.** Correlation clustering of circulating metabolites (**B**) and lipids (**C**) altered by BAT ablation. *n* = 8 for BAT-ablated mice and *n* = 10 for littermate controls. BAT-, BAT-ablated mice; BAT+, littermate controls. C-1h, 6 °C for 1 hour; C-3h, 6 °C for 3 hours; C-3d, 6 °C for 3 days; TN-3d, 28 °C for 3 days. Mice were provided with ad libitum access to food and water throughout the experiment. **D-E.** Representative quantitative profiles of known BAT-regulated metabolites (**D**) and lipids (**E**). *n* = 8 for BAT-ablated mice and *n* = 10 for littermate controls. Statistic: unpaired *t*-test, ^∗^*p* < 0.05; ^∗∗^*p* < 0.01; ^∗∗∗^*p* < 0.001. **F.** Representative quantitative profiles of metabolites and lipids newly identified as BAT-regulated in this study. *n* = 8 for BAT-ablated mice and *n* = 10 for littermate controls. Statistic: unpaired *t*-test, ^∗^*p* < 0.05; ^∗∗^*p* < 0.01; ^∗∗∗^*p* < 0.001.

## RESULTS

### The impact of BAT on circulating metabolome and lipidome *in vivo*

To study how BAT affects circulating metabolite and lipid profiles *in vivo*, we devised the following strategy (**Figure 1A**). First, we generated a BAT ablation mouse model in which peroxisome proliferator-activated receptor-γ (*Pparg*) was deleted in uncoupling protein 1 (UCP1)-expressing adipocytes (*Ucp1*-Cre; *Pparg*^flox/flox^, herein BAT- mice) (Step 1). We found that the presumptive interscapular BAT (iBAT) in BAT-ablated mice was largely replaced by unilocular adipocytes and fibrotic tissue relative to that of littermate controls (*Pparg*^flox/flox^, herein BAT+ mice) (**Figure S1A**). iBAT weight in BAT- mice was reduced compared to BAT+ mice by approximately 4-fold in males and 2-fold in females (**Figure S1B**), while overall body weight remained unchanged on a regular diet under room temperature (**Figure S1C**). Consistently, the expression of BAT marker genes, including *Ucp1*, *Cidea*, *Elovl3*, *Cox7a,* and *Pparg*, was significantly reduced in BAT- mice relative to those in BAT+ mice (**Figure S1D**). These mice were kept at room temperature (RT) and then exposed to cold temperature (6 °C for 1 and 3 hours), chronic cold (6 °C for 3 days), and subsequently acclimated to thermoneutrality (TN, 28 °C for 3 days) under an ad-libitum feeding condition, from which we harvested their serum samples (Step 2). Note that we adapted this protocol to avoid hypothermia in BAT- mice rather than by directly transferring from thermoneutrality to cold. These samples were then analyzed by LC-MS for metabolomics and lipidomics (Step 3). Since the initial data analysis was intended as an exploratory discovery to generate a candidate list while avoiding premature discarding of biologically important candidates, we used unadjusted p-values as an initial filter rather than multiple-comparison correction. Candidates were subsequently prioritized based on their biological consistency across complementary datasets, including tissue enrichment, extracellular fluids, tissue explant and cell-secretion assays, and cross-species human data. As discussed later, we tested the biological function of prioritized candidates.

The analysis identified 206 metabolites and 249 lipid species whose circulating profiles were altered by BAT ablation (**Table S1**). These metabolites and lipids were classified into distinct dynamic patterns (Cluster I to IV for metabolites, Cluster I to VI for lipids), with metabolically related molecules grouped together, and the first three metabolite clusters and the first four lipid patterns accumulated in BAT-ablated mice (Step 4). For example, circulating BCAAs (Val, Leu, Ile) in BAT-ablated mice were elevated relative to those in littermate control mice during cold exposure (Cluster I) **(Figure 1B**). This result is consistent with previous studies showing that BAT actively takes up BCAAs in response to cold exposure, while a defect in BCAA catabolism in BAT leads to elevated circulating BCAA levels ^12,29–31^. Similarly, glutathione-related metabolites, including glutamyl-glutamic acid (γ-Glu-Glu), pyroglutamic acid (Pyro-Glu), S-adenosylmethionine (SAM), and oxidized glutathione (GSSG) were elevated in BAT-ablated mice relative to control mice under thermoneutrality (Clustered III). Among lipids, the larger, more hydrophobic molecules, including lipoprotein-derived lipids, including triglycerides (TGs), diglycerides (DGs), cholesterol esters (CEs), ceramides (CER), and phospholipids, were classified in the first four clusters in which their circulating levels were elevated in BAT-ablated mice relative to control (BAT+) mice. In contrast, smaller, more hydrophilic free lipids, such as acylcarnitines (CAR) and hydroxy fatty acids (OHFA), clustered in the last two groups and were more abundant in the serum of BAT+ mice compared to those in BAT- mice (**Figure 1C**).

The quantitative profiles of these clusters were presented as follows. Circulating BCAA (Val, Leu, and Ile) levels in BAT-ablated mice remained elevated relative to those in control mice during cold exposure (**Figure 1D, Figure S1E**). Notably, BAT ablation reduced glutathione (GSH) levels in response to cold exposure. Conversely, BAT-ablation mice exhibited elevated GSSG levels compared to control mice at thermoneutrality (**Figure S1E**). These results are consistent with our previous study showing that BCAA catabolism in BAT contributes to GSH synthesis and redox homeostasis ^12^. It has been reported that BAT plays a key role in circulating lipid metabolism by decreasing circulating TG levels ^14,32,33^. In agreement, circulating levels of TG (e.g., TG 14:0/16:0/18:1) and DG (e.g., DG 18:2/18:2) in BAT-ablated mice were higher compared to those in control mice during cold exposure (**Figure 1E**).

Beyond these known metabolites and lipids, our analysis uncovered new insights. For instance, BAT-ablated mice showed elevated circulating serotonin levels compared to control mice. This is intriguing, given previous work showing that BAT actively imports serotonin, which acts as a negative regulator of thermogenesis ^34–36^. Moreover, BAT-ablation mice exhibited elevated circulating AMP levels (**Figure 1F**). We also found that several lipid species were significantly lower in the serum of BAT-ablation mice than in control mice in response to cold exposure. These included medium-chain acylcarnitines (e.g., CAR 8:0), hydroxy stearic acid (OHFA 18:0), and dihydroxy octadecadienoic acid (DiHODE) (**Figure 1F, Figure S1E**). On the other hand, there was no difference in serum levels of fatty acids, including palmitic acid, stearic acid, oleic acid, linoleic acid, and arachidonic acid, between the two groups (**Figure S1E**). This apparent difference from previous studies, in which acute cold exposure increased circulating fatty acids under fasted conditions ^37–39^, is likely because the present study was conducted under an ad libitum condition during chronic cold adaptation over three days. One exception was docosahexaenoic acid (DHA), which showed a modest but significant reduction in BAT-ablated mice compared with control mice after cold exposure. Additionally, 12,13-diHOME showed a trend toward reduction in BAT-ablated mice, although it did not reach statistical significance. Together, the comprehensive serum analyses using the BAT-ablation mouse model identified known and unknown metabolites and lipids whose circulating profiles are linked to BAT *in vivo*.

### Tissue origins of the BAT-linked circulating metabolites and lipids

To determine the contributions of BAT and other organs to the BAT-linked metabolites in circulation, we developed the quantitative profiles of these metabolites and lipids across twelve mouse tissues (**Figure 2A**). Subsequently, we examined if any of these metabolites and lipids were detected from i) the conditioned media in *ex vivo* cultured iBAT explants, ii) extracellular fluids isolated from iBAT as well as epididymal WAT as a comparison, and iii) the conditioned media in cultured brown adipocytes.

**Figure 2.**
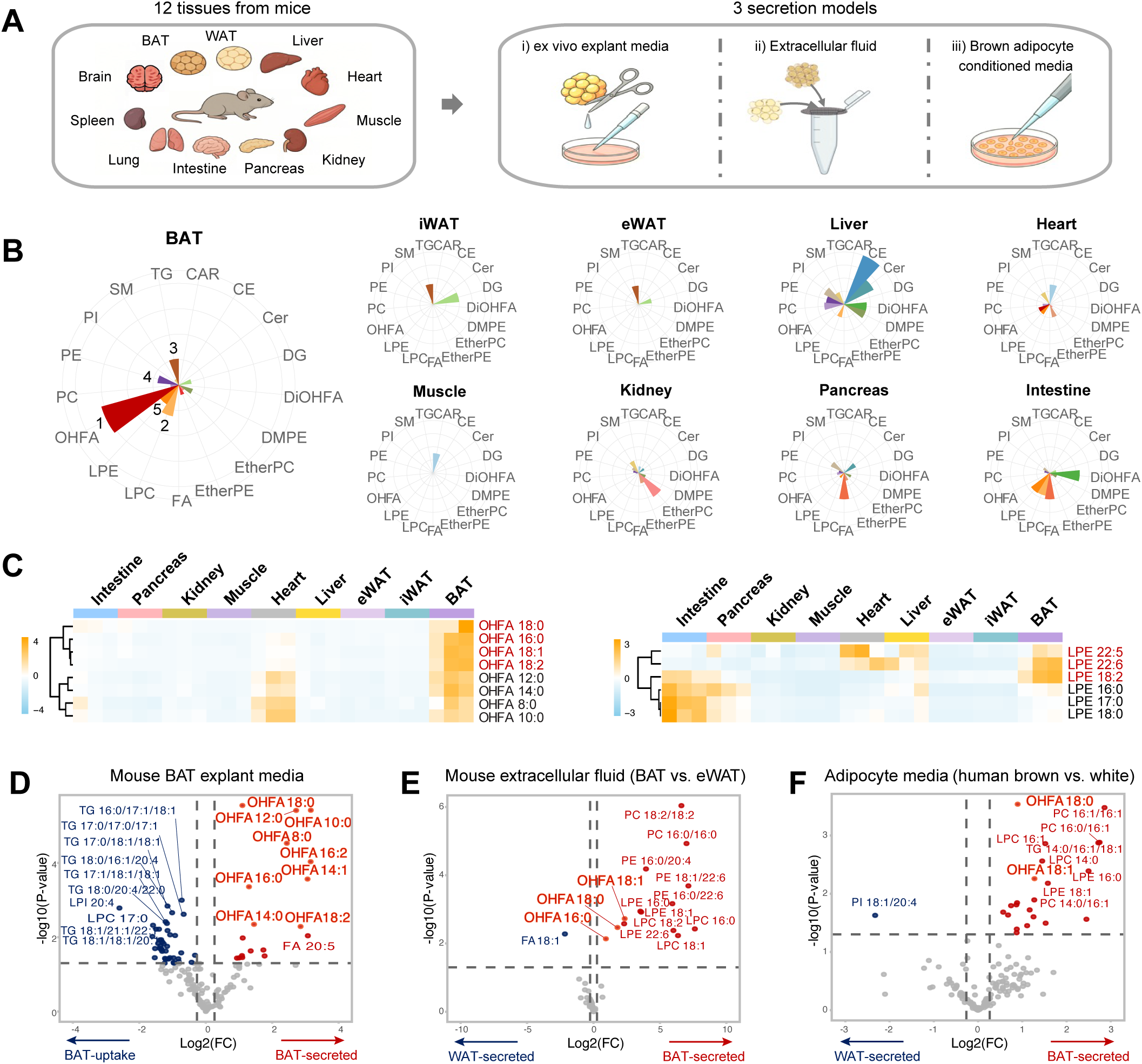
Tissue origins of the BAT-responsive circulating lipids. **A.** Schematic of the source mapping strategy. **B.** Source mapping of the BAT-responsive circulating lipids across nine mouse tissues. Brain, lung, and spleen, which did not show strong lipid enrichment, are not shown here. *n* = 3 for each tissue. **C.** Distribution of carbon chain length and degree of unsaturation of BAT-responsive circulating OHFA and LPE lipid species across nine tissues. Brain, lung, and spleen, which did not show strong lipid enrichment, are not shown here. *n* = 3 per group. **D.** Volcano plot of lipids in 12-hour conditioned media from mouse BAT explants compared with fresh media. *n* = 3 for each group. Statistic: unpaired *t*-test. *p* < 0.05, FC >1.2 or <0.83. Top 10 up- and down-regulated lipids were labeled by name in the plot. **E.** Volcano plot of lipids in extracellular fluids from mouse brown and epididymal white adipose tissues. *n* = 3 for each group. Statistic: unpaired *t*-test. *p* < 0.05, FC >1.2 or <0.83. **F.** Volcano plot of lipids in 12-hour conditioned media from differentiated human brown and white adipocytes. *n* = 3 for each group. Statistic: unpaired *t*-test. *p* < 0.05, FC >1.2 or <0.83. Top 10 up-regulated lipids were name-labeled in the plot. The dot and label of hydroxy fatty acids were outlined in orange.

The analyses made two observations. First, most of the BAT-linked metabolites were detected in multiple tissues other than BAT (**Figure S2A**). For example, BAT actively takes up BCAAs upon cold exposure; however, its role in secreting BCAAs into circulation is minimal. This is consistent with a recent study that analyzed the metabolomic profiles of arteriovenous serum across BAT ^40^. Similarly, BCAA-derived metabolites, such as GSH, contribute to the circulating pool, whereas the liver predominantly produces GSH at the highest levels ^41–43^. Second, we found that many BAT-linked lipids, especially hydroxy fatty acids (OHFAs), were highly enriched in BAT, alongside other lipids such as lysophosphatidylcholines (LPC), TGs, phosphatidylethanolamines (PE), and lysophosphatidylethanolamines (LPE) (**Figure 2B**). A similar enrichment pattern was observed when analyzing the entire tissue lipid class beyond BAT-linked circulating lipids (**Figure S2B**). Although some tissues shared enrichment of the same lipid class, the distribution of carbon chain lengths and degrees of saturation differed markedly: the heart contained high levels of OHFAs, but these were predominantly medium-chain OHFAs, such as C8:0 and C10:0. On the other hand, long-chain OHFAs, especially C18, were uniquely enriched in BAT relative to other tissues (**Figure 2C**). The intestine was enriched in saturated LPEs, whereas BAT contained highly unsaturated species, including C18:2, C22:5, and C22:6 (**Figure 2C**). Likewise, PEs enriched in BAT tended to have one monounsaturated chain, such as C16:1 and C18:1 (**Figure S2C**).

To verify if these are BAT-derived circulating lipids, we next harvested conditioned media from ex vivo cultured BAT explants. We found that OHFAs (e.g., 18:0, 10:0, and 12:0) were abundantly secreted from ex vivo cultured BAT explants, whereas TGs (e.g., 18:0/16:1/20:4, 16:0/17:1/18:1, and 17:0/18:1/18:1) in the media were taken up by BAT explants following 12 hours of culture (**Figure 2D**). Next, we analyzed lipids in extracellular fluid (EF) derived from BAT and epididymal white adipose tissue (eWAT), as previously reported ^12^. We found that lipids in eWAT-derived extracellular fluids contained largely triglycerides, whereas phospholipids were more abundant in BAT-derived extracellular fluids, including PC and PE. Notably, OHFA 16:0, 18:0, and 18:1 were significantly enriched in BAT-derived EF relative to eWAT-derived EF (**Figure 2E**). Since BAT is composed of many cell types, including progenitor cells and immune cells, we next asked whether any of these lipids are derived from differentiated brown adipocytes. To this end, we differentiated human brown adipocytes and white adipocytes as a comparison and collected their conditioned media. We identified several lipid species that were actively released from brown adipocytes into the media relative to white adipocytes, such as PCs (e.g., C16:1/C16:1 and C16:0/C16:1), LPE (C16:0, C18:1), and OHFAs (C18:0 and C18:1) (**Figure 2F**). Collectively, the analyses found that hydroxy fatty acids, which showed a marked reduction in circulation in BAT-ablated mice, were highly enriched in brown adipose tissue and secretory media, supporting their potential identity as BAT-derived lipids.

### Cross-species integration of BAT-linked lipids in mouse and human

We next investigated whether any of the above-identified BAT-derived circulating lipids were also associated with BAT activity in humans. To this end, serum samples were collected from healthy young adults (*n* = 84; male, 65; female, 19; average age, 23.2 years) at 27°C and after 1 hour of mild cold exposure at 19°C under a fasted condition (**Figure 3A, Figure S3A, Table S2**). One hour after blood sample collection (2 hours after c old exposure), we used the standardized uptake value (SUV) in positron emission tomography-computed tomography with 18F-fluorodeoxyglucose (^18^FDG-PET) to determine the BAT activities of the subjects ^44^. Subjects were classified equally into the high BAT group (SUV≥3.4, *n* = 42) and the low BAT group (SUV<3.4, *n* = 42) based on the median of SUV (**Figure 3B**). For subject classification, we used median-split dichotomization, an unbiased statistical method widely used in ^18^FDG-PET-based BAT studies ^29,45–49^. The cut-off value of SUV≥3.4 is aligned with a previous study that used SUV>4.0 ^29^. We first analyzed the hydrophilic lipids using a methanol-based extraction procedure coupled with an LC-MS method employing acetonitrile as the elution solvent. We also extended the analyses to hydrophobic lipid species using a chloroform-based extraction protocol coupled with an LC-MS method utilizing isopropanol for elution.

**Figure 3.**
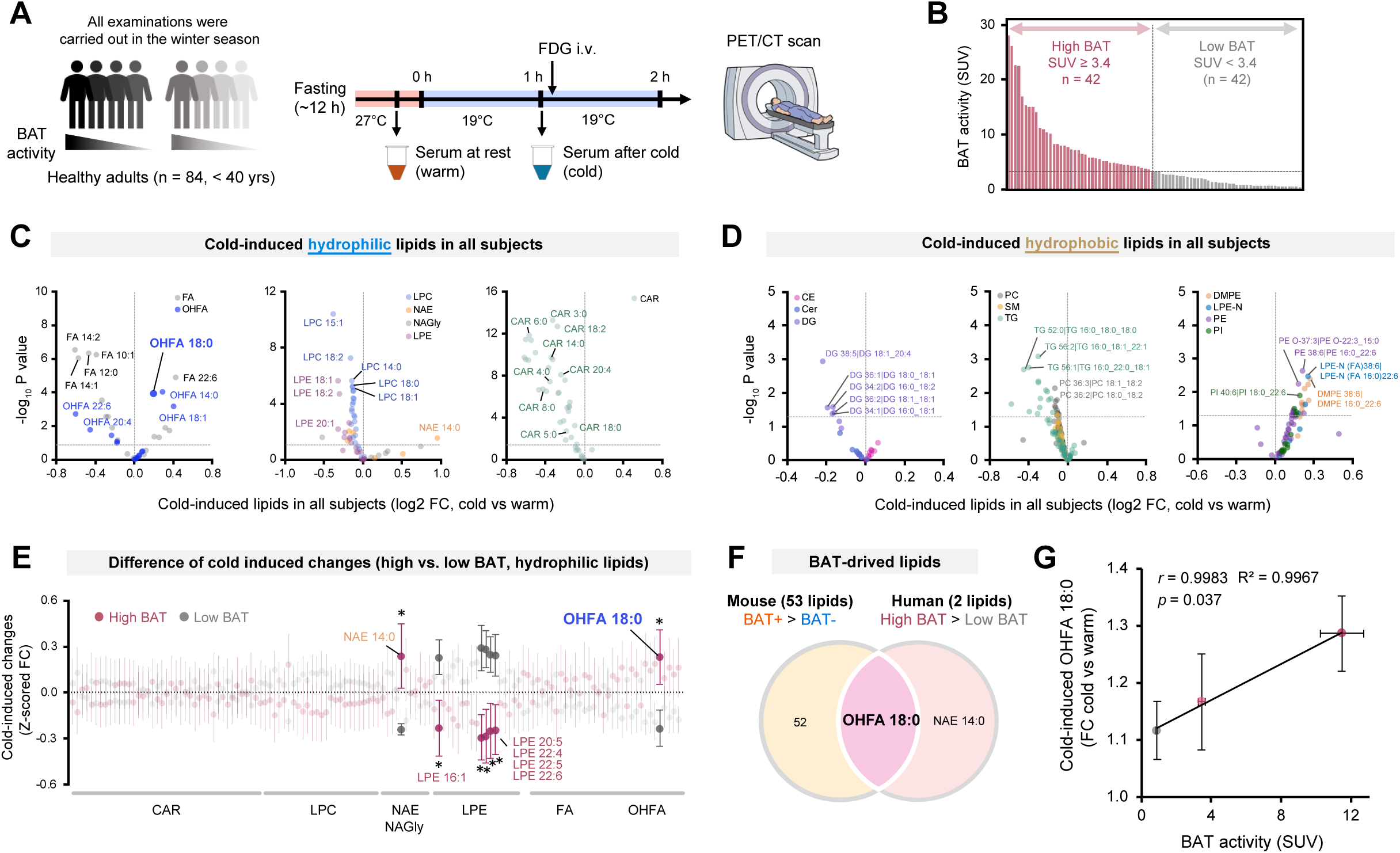
Cross-species integration of BAT-linked lipids in mouse and human. **A.** Sample information of the human cohort. The study included 84 healthy adults (mean age 23.2 years, 65 males and 19 females) with varying BAT activities. Serum samples were collected from these subjects at 27 °C (herein warm) and after 1 hour of mild cold exposure at 19 °C (herein cold). **B.** A total of 84 healthy adults with varying BAT activities were separated into high and low BAT groups, using the median SUV value of 3.4 as the cutoff. High BAT, SUV ≥ 3.4, *n* = 42; low BAT, SUV < 3.4, *n* = 42. **C.** Volcano plot of cold-induced changes in hydrophilic lipids among all subjects. *n* = 84 for each group (cold vs. warm). Statistic: unpaired *t*-test. *p* < 0.05. **D.** Volcano plot of cold-induced changes in hydrophobic lipids among all subjects. *n* = 80 for each group (cold vs. warm). Statistic: unpaired *t*-test. *p* < 0.05. **E.** Difference of cold-induced changes in hydrophilic lipids between the high BAT and low BAT groups. Cold-induced changes, Z-scored FC (cold/warm). *n* = 42 for each group (high BAT vs. low BAT). Statistic: unpaired *t*-test. ^∗^*p* < 0.05. **F.** Overlap analysis of lipids elevated in high BAT humans versus low BAT humans and those elevated in control mice with intact BAT versus BAT-ablated mice. **G.** Correlation between FDG uptake in BAT and cold-induced fold change (FC) in circulating OHFA 18:0. Statistic: correlation analysis and coefficient of determination (*R*^2^). Participants were classified into three groups according to the tertile: high BAT (*n* = 28, SUV ≧ 5.2), middle BAT (*n* = 28, 2.0 ≦ SUV < 5.2), and low BAT groups (*n* = 28, SUV < 2.0).

Among hydrophilic lipids, we found that cold exposure significantly elevated serum levels of OHFAs (14:0, 18:0, and 18:1) as well as FA 22:6 (**Figure 3C**). On the other hand, serum levels of fatty acids (FA10:1, 12:0, 14:1, and 14:2) were reduced by cold exposure. We also found that cold exposure significantly reduced serum levels of LPE (18:1, 18:2, 20:1) and LPC (15:1, 18:2, 14:0, 18:0, and 18:1), whereas cold modestly elevated N-acylethanolamine (NAE 14:0) levels. Similarly, serum levels of acylcarnitines (CAR 3:0, 4:0, 5:0, 6:0, 8:0, 14:0, 18:0, 18:2, 20:4) decreased following cold exposure. For hydrophobic lipids, serum levels of DGs (e.g., 18:1/20:4, 18:1/18:1, and 16:0/18:2), TGs (16:0/18:0/18:0, 16:0/18:1/22:1, and 16:0/22:0/18:1), and PCs (18:1/18:2 and 18:0/18:2) were reduced following cold exposure. In contrast, PE (16:0/22:6), PI (18:0/22:6), LPE-N (16:0/22:6), and DMPE (16:0/22:6) were elevated under cold exposure (**Figure 3D**). The list of lipids and their serum levels following cold exposure is provided in **Table S3.** Subsequently, we asked the extent to which temperature-dependent changes in these lipid species were linked to BAT activity. To this end, we analyzed cold-induced changes in serum lipids in the high-BAT group compared with the low-BAT group (**Figure 3E, Figure S3B**). We found that cold-induced elevation of OHFA 18:0 was significant in the high BAT group, but not in the low BAT group. Similarly, NAE 14:0 levels increased selectively in the high BAT group following cold exposure. On the other hand, serum levels of LPEs (20:5, 22:4, 22:5, 22:6) showed a significant decrease by cold exposure in the high BAT group, but this reduction was not seen in the low BAT group (**Figure 3E, Table S4**).

To integrate all lipid datasets, we compared circulating lipids associated with BAT activity in humans with those identified in mice. Although 53 serum lipid species were reduced in BAT-deficient mice relative to BAT-intact controls, OHFA 18:0 emerged as a common BAT-linked lipid across both species (**Figure 3F**). We then performed a subgroup analysis based on tertiles (low, mid, and high, with *n* = 28 per group) to ensure equal sample sizes within each group (**Figure S3C**). We found a significant positive correlation between cold-induced OHFA 18:0 and BAT activity (**Figure 3G**). On the other hand, no significant correlation was observed with other candidate lipids, although the initial screening identified LPE 22:6 as BAT-linked lipids in mice and humans (**Figure S3D, E)**. In addition to the tertile-based analyses, there was a modest but significant correlation between cold-induced OHFA 18:0 and BAT SUV activity in subjects with SUV≥ 1 (**Figure S3F, G**). We note substantial interindividual variation in cold-induced OHFA 18:0 in individuals with SUV<1, suggesting the possibility that other tissues may compensate for circulating 3-OHSA in individuals with very low BAT activity. Overall conservation in lipid profiles between mouse and human serum signatures was limited, possibly reflecting species-specific differences in lipid metabolism, cold-exposure protocol, diet, and body size, as well as differences in the contribution of shivering vs. non-shivering thermogenesis to cold adaptation. Nonetheless, OHFA 18:0 was consistently elevated by cold exposure in both mice and humans in a BAT-dependent manner, representing the circulating lipid species associated with cold-induced BAT activity across species.

### 3-OHSA is a cold-inducible BAT-derived lipid released into circulation

Given the unique profile of OHFA 18:0 among BAT-derived lipids, we next aimed to determine the location of the hydroxyl group in OHFAs. To this end, we compared the retention times and MS/MS spectra of OHFA 18:0 with authentic standards (2-hydroxy stearic acid, 3-hydroxy stearic acid, 9-hydroxy stearic acid, and 12-hydroxy stearic acid). This analysis revealed a perfect match with 3-hydroxy stearic acid (3-OHSA or 3-OHFA 18:0) (**Figure 4A, Figure S4A**). All the OHFAs detected in our lipidomics exhibited the diagnostic MS/MS feature of 3-hydroxy fatty acids (3-OHFAs) ^50^ (**Figure S4B**).

**Figure 4.**
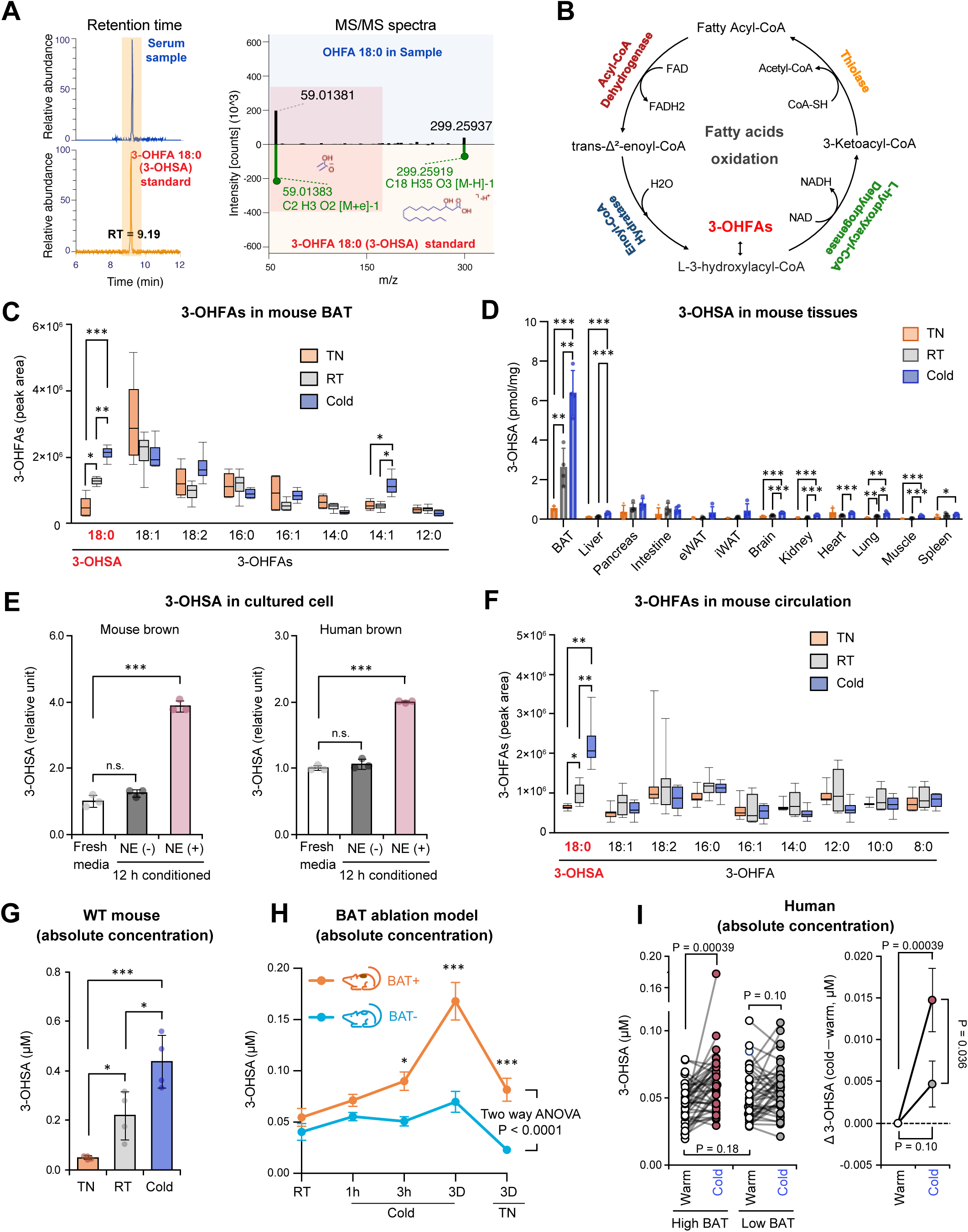
3-OHSA is a cold-inducible BAT-derived lipid released into circulation **A.** Determination of the hydroxyl group position in hydroxy stearic acid by matching retention time and MS/MS spectra with a chemically synthesized standard. 3-OHSA, 3-hydroxy stearic acid. **B.** Schematic diagram of fatty acid oxidation. The enzymes involved in the four steps of fatty acid oxidation are highlighted in four different colors. **C.** Relative levels of 3-hydroxy fatty acids in BAT under TN (2 weeks), RT, and cold (3 days). *n* = 4 per group. Data are shown as box with whiskers min to max. **D.** Absolute concentration of 3-OHSA in the indicated tissues of mice under TN (4 weeks), RT, and cold (3 days). *n* = 4 per group. Data are shown as mean ± SD. **E.** Quantitative levels of 3-OHSA in fresh and 12 hour-conditioned media from mouse and human brown adipocytes, with or without norepinephrine stimulation. *n* = 3 per group. Data are shown as mean ± SD. **F.** Relative levels of circulating 3-hydroxy fatty acids under TN (2 weeks), RT, and cold (3 days). *n* = 6 per group. Data are shown as box with whiskers min to max. **G.** Absolute concentration of 3-OHSA in serum from mice exposed to TN (4 weeks), RT, and cold (3 days). *n* = 4 per group. Data are shown as mean ± SD **H.** Absolute concentrations of 3-OHSA in BAT-ablated mice and littermate controls during adaption to cold (1hour, 3 hours, and 3days) and to TN (3days) under ad libitum conditions. *n* = 8 for BAT-ablated mice and *n* = 10 for controls. Data are shown as mean ± SEM. **I.** Serum levels of 3-OHSA in healthy adult humans under warm (27°C) and cold (19°C) conditions. Left panel: Absolute serum concentrations of 3-OHSA in humans with high or low BAT. Right panel: Cold-induced changes in circulating 3-OHSA in humans with high or low BAT. High BAT, SUV ≥ 3.4; low BAT, SUV < 3.4. *n* = 42 per group. Statistic (**C**, **D**, **E**, and **G**): unpaired *t*-test; **F**: paired *t*-test; **H**: two-way ANOVA with Šídák’s correct for multiple comparisons; **I**: unpaired *t*-test for high versus low BAT; paired *t*-test for warm versus cold. *p* < 0.05; ^∗∗^*p* < 0.01; ^∗∗∗^*p* < 0.001.

3-OHFAs are generated through active fatty acid oxidation (FAO) in the mitochondria (**Figure 4B**). In response to cold, stearic acid (C18:0) is activated to stearoyl-CoA and imported into the mitochondrial matrix via the carnitine shuttle. Each β-oxidation cycle proceeds by the actions of (i) acyl-CoA dehydrogenases (e.g., ACADL) to trans-2-enoyl-CoA, (ii) enoyl-CoA hydratases (e.g., ECH) to form L-3-hydroxystearoyl-CoA (the immediate 3-OH intermediate), (iii) NAD⁺-dependent oxidation by the L-3-hydroxyacyl-CoA dehydrogenases (e.g., HADH) to 3-ketoacyl-CoA, and (iv) thiolases (e.g., ACAA2, HADHB) to shorten the acyl-chain. Under high FAO flux upon cold exposure, the dehydrogenase step can become transiently rate-limiting (e.g., local NAD⁺/NADH or CoA constraints), resulting in the accumulation of L-3-hydroxystearoyl-CoA. This pool can be diverted by mitochondrial acyl-CoA thioesterases to release the free acid 3-hydroxystearic acid (3-OHSA), or transesterified to 3-hydroxyacylcarnitines for export. In alignment, mRNA expression levels of all four FAO enzymes in BAT were at the highest level compared with those in nine other tissues ^51^ (**Figure S4C**). Quantitative measurements of 3-hydroxy fatty acids across twelve tissues found that BAT contained the highest amounts of 3-OHFAs, including 3-OHFA 18:0, particularly after cold exposure, even though the heart also accumulated some of 3-OHFAs (e.g., medium-chain C8-C12) at room temperature and thermoneutrality (**Figure S4D**).

The BAT content of 3-OHFA 18:0 (3-hydroxy stearic acid, 3-OHSA) was the second most abundant 3-OHFA after 3-OHFA 18:1 at RT; however, it increased significantly following cold exposure, and decreased when acclimated to thermoneutrality (**Figure 4C**). Next, we generated a standard curve using a chemically synthesized standard and quantified the absolute concentration of 3-OHSA in tissues and serum. Notably, BAT exhibited the highest tissue concentration, approximately 20-fold greater than that in other metabolic organs examined, and displayed the strongest temperature-dependent upregulation (**Figure 4D**). The cold-inducible change in BAT is mediated by norepinephrine (NE) signaling, given the significant increase in the release of 3-OHSA into the conditioned media following NE treatment in both differentiated mouse and human brown adipocytes (**Figure 4E**). In circulation, 3-OHSA was the most abundant 3-OHFA species: after 3 days of cold exposure, serum 3-OHSA levels were significantly higher than those in mice acclimated to thermoneutrality, while other OHFA species did not show this pattern under a fed condition (**Figure 4F**). Absolute quantification in mouse serum found that 3-OHSA concentration reached approximately 0.5 µM after 3 days of cold exposure under a fed condition (**Figure 4G**). We note that serum 3-OHSA is unstable after freeze-thaw cycles and, as a result, the absolute concentrations of 3-OHSA in frozen serum samples were lower than those quantified in freshly prepared samples. Nonetheless, analyses of independent cohorts consistently showed that serum 3-OHSA levels increased following 3 hours or longer of cold exposure, but this induction was blunted in BAT- mice (**Figure 4H, Figure S4E**). Serum 3-OHSA levels decreased when mice were acclimated to thermoneutrality for 3 days, and further decreased to a basal level after one week (**Figure S4E**). After one week of acclimation to thermoneutrality, there was no difference in serum 3-OHSA levels between BAT+ vs. BAT- mice. In human serum, the absolute quantification of 3-OHSA showed that cold exposure significantly elevated 3-OHSA concentrations in individuals with high BAT, whereas the cold-inducible change was not observed in individuals with low BAT (**Figure 4I**). Consistent with the results in mice, there was no difference in the basal 3-OHSA concentrations under warm conditions between the two groups. These results suggest that 3-OHSA is a cold-inducible lipid whose induction is linked to BAT activity, while its basal levels at TN did not distinguish the subjects with high vs. low BAT activity.

### 3-OHSA mitochondrial H_2_O_2_ production in the liver

The next key question is whether 3-OHSA is merely a circulating readout of cold-induced BAT activity or has any biological action. Since 3-OHSA is produced as a byproduct of active FAO rather than via a specific enzymatic pathway, targeting ACADL or ECH would disrupt the entire FAO system. As an alternative approach, we examined the extent to which BAT ablation affects liver function under room temperature and cold conditions. We found that BAT ablation led to significantly increased oxidative markers in the liver, including lipid peroxidation, as measured by malondialdehyde (MDA), and protein oxidation, as measured by protein carbonyl content (**Figure 5A**). Next, we asked whether BAT-derived 3-OHSA is a part of BAT-derived mediators that regulate hepatic oxidative stress. Intriguingly, the addition of 3-OHSA to mitochondria isolated from the liver significantly reduced mitochondrial H_2_O_2_ production rate, to an extent comparable to that of FCCP (**Figure 5B**). Of note, this reduction occurred in a dose-dependent manner, with a half-maximal effective concentration (EC_50_) of 0.51 µM, which was within the physiological circulating concentration range (**Figure 5C**). We then assessed the tissue-specificity of 3-OHSA’s action in reducing the H_2_O_2_ production rate by examining the effect of 3-OHSA in metabolic organs, including the liver, skeletal muscle, kidney, and BAT (**Figure 5D**). We found that 3-OHSA significantly reduced mitochondrial H_2_O_2_ production rate relative to vehicle control in the liver and kidney (**Figure 5E**). This effect was not due to changes in hepatic mitochondrial contents (**Figure S5A**). 3-OHSA also decreased mitochondrial H_2_O_2_ production rate in muscle, albeit to a lesser extent than in the liver, whereas no effect was observed on BAT mitochondria. Additionally, 3-OHSA increased mitochondrial respiration; however, the concentration required to affect mitochondrial respiration was much higher than that required for lowering mitochondrial H_2_O_2_ production, with an EC_50_ of ∼5.5 µM **(Figure S5B-D**). These results suggest that, although 3-OHSA at supraphysiological concentrations can increase mitochondrial respiration as other fatty acids do, its physiological role is rather to reduce mitochondrial ROS production.

**Figure 5.**
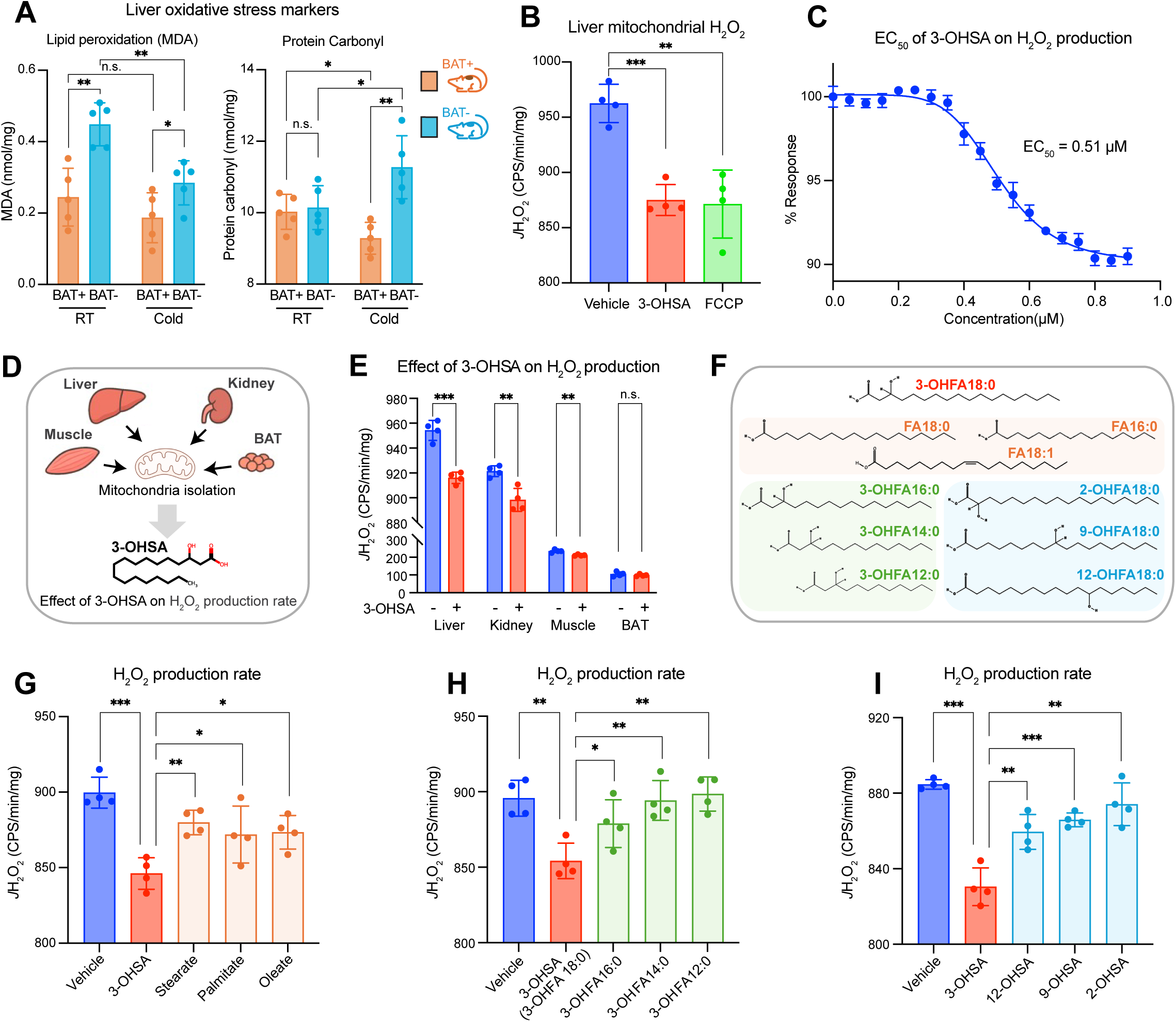
3-OHSA reduces mitochondrial ROS production in the liver. **A.** Hepatic oxidative stress markers in BAT-ablated mice and littermate controls exposed to RT or Cold (3 days). Left, lipid peroxidation marker malondialdehyde (MDA); right, protein oxidation marker carbonyl content. *n* = 5 per group. **B.** H_2_O_2_ production rate of liver mitochondria stimulated with vehicle (ethanol), 3-hydroxy stearic acid (3-OHSA), or FCCP. *n* = 4. **C.** EC_50_ of 3-OHSA on liver mitochondrial H_2_O_2_ generation rate. *n* = 4. Data are shown as mean ± SEM. **D.** Schematic overview of the approach used to investigate the effect of 3-OHSA on H_2_O_2_ production across metabolic organs. **E.** Effect of 3-OHSA on the H_2_O_2_ generation rate in isolated mitochondria from the indicated organs. 3-OHSA concentration, 0.5 μM. *n* = 4. **F.** The functional specificity of 3-OHSA and other lipid species on liver mitochondrial H_2_O_2_ generation. **G.** Quantification of liver mitochondrial H_2_O_2_ generation rates in response to vehicle, 3-OHSA, or non-hydroxylated fatty acids at 0.5 μM. *n* = 4. **H.** Quantification of liver mitochondrial H_2_O_2_ generation rates in response to vehicle, 3-OHSA, or the indicated 3-hydroxy fatty acids at 0.5 μM. *n* = 4. **I.** Quantification of liver mitochondrial H_2_O_2_ generation rates in response to vehicle, 3-OHSA, or hydroxy fatty acids with varying hydroxyl position at 0.5 μM. *n* = 4. Statistic (**A**, **B**, **E**, **G**, **H**, and **I**): unpaired *t*-test, *p* < 0.05; ^∗∗^*p* < 0.01; ^∗∗∗^*p* < 0.001. Data are shown as mean ± SD.

Next, we asked to what extent this reduction in mitochondrial H_2_O_2_ production was unique to 3-OHSA. To this end, we compared its activity with three classes of controls: non-hydroxylated fatty acids (stearic acid, palmitic acid, oleic acid), 3-hydroxy fatty acids with varying carbon chain lengths (12:0, 14:0, 16:0), and hydroxy fatty acids with varying hydroxyl position (2-OHSA, 12-OHSA, 9-OHSA) (**Figure 5F**). These lipid species were independent of BAT activity as BAT ablation did not affect their circulating levels *in vivo*. Given its EC_50_ and the physiological range observed in circulation, we performed the assays using 0.5 µM, along with 3-OHSA as a comparison, and observed the following: First, 3-OHSA exerted a significantly stronger H_2_O_2_-lowering effect than non-hydroxylated fatty acids (**Figure 5G**). Second, the magnitude of the effect depended on fatty acid chain length, following the order 3-OHFA 18:0 (3-OHSA) > 16:0 > 14:0 > 12:0 (**Figure 5H**). Lastly, the position of the hydroxyl group was critical, with 3-OHSA showing a stronger effect than 12-OHSA, 9-OHSA, or 2-OHSA (**Figure 5I**). These results suggest that 3-OHSA reduces mitochondrial H_2_O_2_ production in the liver in a manner dependent on both the presence and position of the hydroxy group, as well as fatty-acid chain length.

### 3-OHSA reduces liver mitochondrial membrane potential and oxidative stress

We next examined the underlying mechanism by which 3-OHSA reduces hepatic ROS production. Since mitochondrial ROS production is often accompanied by reduced mitochondrial depolarization ^52–55^, we tested whether 3-OHSA alters mitochondrial membrane potential (ΔΨm). To this end, we isolated mitochondria from the liver and measured the rate of ΔΨm polarization using safranine in the presence of malate, glutamate, pyruvate, and succinate. Note that a decrease in safranine intensity represents a polarized membrane (a signal decline in response to the addition of mitochondria in **Figure 6A**). As expected, the chemical uncoupler FCCP markedly reduced the rate of Δψm polarization. To our surprise, 3-OHSA similarly reduced the rate of ΔΨm polarization to a degree comparable to that of FCCP (**Figure 6A**). This effect was dose-dependent, with an EC_50_ of 0.57 µM (**Figure 6B**). The dose-response relationship of 3-OHSA-induced ΔΨm depolarization closely mirrored its effect on mitochondrial H_2_O_2_ production, and within the physiological range of 3-OHSA in circulation. We then assessed the specificity of 3-OHSA to ΔΨm polarization rate by comparing the effect with those of non-hydroxylated fatty acids (stearic acid, palmitic acid, oleic acid), 3-hydroxy fatty acids with short carbon chain lengths (12:0, 14:0), and a hydroxy fatty acid with an alternative hydroxyl position (2-OHSA). Similar to the effect on mitochondrial H_2_O_2_ production, 3-OHSA at 0.5 µM was more potent than other lipid species tested at the same concentration, although some of them, such as oleate and palmitate, also reduced ΔΨm, albeit with lesser potency (**Figure 6C**). These results suggest that the position of the hydroxyl group and fatty acid chain length are critical for the reduction effect of 3-OHFA on ΔΨm polarization. Additionally, 3-OHSA increased state 4 mitochondrial respiration and reduced the ATP synthesis rate, leading to a reduced P/O ratio, although this effect required a higher dose (**Figure S6A**).

**Figure 6.**
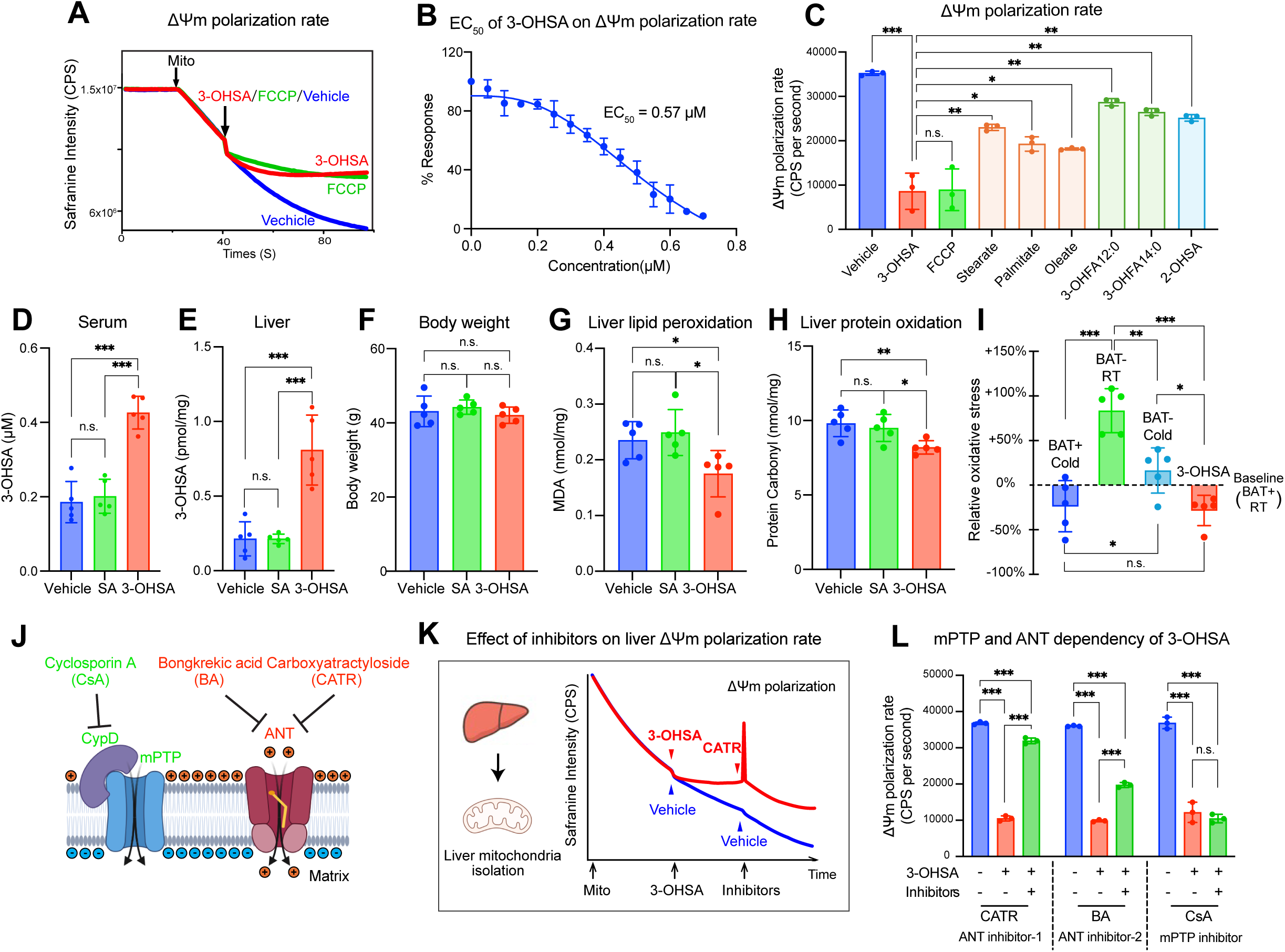
3-OHSA reduces hepatic mitochondrial membrane potential and oxidative stress. **A.** Representative traces of hepatic mitochondrial membrane potential (ΔΨm) polarization in response to vehicle, 3-OHSA, or FCCP. **B.** EC_50_ of 3-OHSA on hepatic ΔΨm polarization rate. *n* = 3 per group. **C.** The action specificity of the 3-OHSA on the hepatic ΔΨm polarization rate. *n* = 3 per group. **D-E.** Absolute concentration of 3-OHSA in serum (**D**) and liver (**E**) from HFD-induced obese (DIO) mice supplemented with vehicle, stearic acid, or 3-OHSA for two weeks. *n* = 5 per group. **F.** Body weight of mice supplemented with vehicle, stearic acid, or 3-OHSA for two weeks. *n* = 5 per group. **G.** Abundance of lipid oxidative stress marker malondialdehyde (MDA) in the liver of mice supplemented with vehicle, stearic acid, or 3-OHSA for two weeks. *n* = 5 per group. **H.** Levels of protein oxidation marker carbonyl in the liver of mice supplemented with vehicle, stearic acid, or 3-OHSA for two weeks. *n* = 5 per group. **I.** Percentage change in liver oxidative stress markers (MDA) in the indicated groups, with BAT+ mice under RT as the baseline. *n* = 5 per group. **J.** Schematic of inhibitors targeting the potential mediator channels of 3-OHSA action. **K.** Representative traces of hepatic ΔΨm polarization in response to 3-OHSA, CATR, or vehicle (indicated by arrow heads). **L.** Effects of ANT inhibitors BA (20 µM) and CATR (1 µM), or mPTP inhibitor CsA (1 µM), on 3-OHSA-induced liver ΔΨm polarization rate. *n* = 3. Statistic (**C**-**I** and **L**): unpaired *t*-test, *p* < 0.05; ^∗∗^*p* < 0.01; ^∗∗∗^*p* < 0.001. Data are shown as mean ± SD.

To examine the metabolic consequences *in vivo*, we next administered BSA-conjugated 3-OHSA intraperitoneally to HFD-induced obese (DIO) mice for two weeks at a dose of 8 mg/kg/day. As controls, we used BSA-conjugated stearate and unconjugated BSA (vehicle). First, we validated using the absolute quantification method that 3-OHSA treatment increased circulating 3-OHSA levels to 0.43 µM ± 0.04, which were comparable to those observed following cold exposure (**Figure 6D**). We also found that 3-OHSA administration resulted in approximately a 3-fold increase in the liver, slightly higher but within the physiological range (**Figure 6E**). On the other hand, this induction was not observed in the heart or skeletal muscle, although a slight increase was found in the kidney (**Figure S6B**). After 2 weeks of treatment, we found no differences in body weight among mice treated with vehicle, 3-OHSA, or stearate (**Figure 6F**). We also did not find any difference in fat mass, lean mass, and tissue weight between the treatment groups (**Figure S6C-E**). However, 3-OHSA reduced hepatic oxidative stress in DIO mice: the liver from 3-OHSA-treated mice exhibited significantly lower levels of oxidative stress markers, including lipid peroxidation (MDA) (**Figure 6G**) and protein oxidation (**Figure 6H**), compared with those in vehicle-treated mice. On the other hand, no change in hepatic redox state was observed with stearate treatment at the same dose. We then calculated the relative change in hepatic oxidative marker MDA in response to BAT ablation, cold exposure, and 3-OHSA treatment compared to those in BAT+ mice at RT as a reference in each experiment. The analysis showed that cold exposure in BAT+ (control) mice significantly reduced hepatic oxidative stress by 24%, whereas BAT-ablation increased oxidative stress by 83% at RT. The cold-induced reduction in hepatic oxidative stress was significantly abrogated in BAT-less mice. Importantly, 3-OHSA treatment markedly reduced oxidative stress by 28%, nearly equivalent to what we observed after cold exposure (**Figure 6I**).

Lastly, we examined how 3-OHSA reduced the hepatic mitochondrial membrane potential. Adenine nucleotide translocase (ANT) has been implicated in modulating mitochondrial membrane potential, thereby reducing ROS production ^56–58^. Among the three forms of ANT (ANT1, 2, 3), ANT2 is the dominant form that is highly expressed in the liver and kidney (**Figure S6F**). Given that the liver is a major target of 3-OHSA, we tested the extent to which ANT2 mediates 3-OHSA’s effect on ΔΨm polarization rate in the liver. Alternatively, we tested whether 3-OHSA acts through the mitochondrial permeability transition pore (mPTP), which is regulated by Cyclophilin D (CypD) and facilitates proton leak ^59–61^. To this end, we acutely inhibited mPTP using the CypD inhibitor cyclosporin A (CsA) and blocked ANT with its inhibitors, carboxyatractyloside (CATR) and bongkrekic acid (BA) (**Figure 6J**). Addition of mitochondrial substrates (malate, pyruvate, glutamate, and succinate) to liver-derived mitochondria leads to ΔΨm polarization, whereas addition of 3-OHSA at 0.5 µM potently reduces the rate of polarization (**Figure 6K**). It is notable that this effect was reversible, as the addition of CATR restored ΔΨm (note: the transient peak immediately after CATR addition is an artifact caused by the injection). The results indicate that 3-OHSA’s action on hepatic mitochondria is not a toxic nonspecific effect; rather, it exerts a specific biological action to modulate ΔΨm. A dose titration study further showed that CATR at lower doses (0.1-5 µM) blocks the effect of 3-OHSA on H_2_O_2_ production in a dose-dependent manner (**Figure S6G**), even though CATR at 1 µM did not affect basal mitochondrial H_2_O_2_ production in the absence of 3-OHSA (**Figure S6H**). Consistently, addition of bongkrekic acid (BA) significantly, though partly, reversed the 3-OHSA-induced reduction in ΔΨm polarization rate (**Figure 6L**). In contrast, CsA did not restore the effect of 3-OHSA. Subsequently, we tested if other lipids, such as 2-OHSA and 3OHFA12:0, have a similar mechanism; however, ANT inhibition with CATR or BA did not alter the effects of 2-OHSA and 3OHFA12:0 on reducing ΔΨm (**Figure S6I, J**), indicating that these lipids act on mitochondria independently of ANT2. As a complementary approach, we acutely depleted ANT2 using siRNA in primary hepatocytes (**Figure S6K)**. Consistent with the pharmacological inhibition, siRNA-mediated knockdown of ANT2 partially but significantly attenuated the 3-OHSA-induced effects on ΔΨm (**Figure S6L**). These findings suggest that 3-OHSA has a unique action in lowering hepatic ΔΨm and oxidative stress, and that this action is mediated, at least in part, through ANT2, with additional mechanisms likely involved.

## DISCUSSION

Historically, BAT-derived factors (batokines) have been searched primarily through analyses of conditioned media from cultured brown/beige adipocytes or tissue explants. In contrast, the present study employed a distinct approach by comprehensively analyzing serum metabolome and lipidome from BAT-ablated mice to determine the impact of BAT on the circulating landscape *in vivo*. Although this approach does not capture paracrine/autocrine factors with minor contributions to circulation, this provides a broadly accessible resource of BAT-associated circulating metabolites, encompassing both candidate BAT-secreted molecules and metabolites systemically regulated through BAT-dependent clearance. Consistent with prior studies demonstrating that BAT promotes BCAA disposal and generates nitrogen-containing mediators ^12,29–31^, genetic BAT ablation resulted in the accumulation of circulating BCAAs and reduced GSH. Our findings also support a role for BAT in circulating lipid remodeling and TG clearance ^32,33^, even though only a subset of lipid alterations was conserved between mice and humans. Conversely, several lipid species found in low-BAT subjects have been linked to adverse metabolic outcomes. For example, LPEs have been suggested as a biomarker for cardiovascular risks in individuals with coronary artery disease, although their causal role remains uncertain ^62,63^. Ceramides are known to impair insulin signaling, suppress lipolysis, and induce lipotoxicity and ER stress ^64–67^, and ceramides containing very long-chain fatty acids (e.g., C22:0, C24:0) were positively associated with cardiovascular disease and myocardial ischemia ^68,69^. In mice, we found that serum ceramide levels (C22:0, C23:0, and C24:0) were elevated in BAT-ablated mice. Collectively, these findings support a model in which active BAT reshapes the circulating lipid and metabolite milieu toward a profile associated with improved cardiometabolic health.

Bidirectional crosstalk between BAT and metabolic organs enables coordinated adaptation to metabolic stressors. During vigorous exercise, skeletal muscle releases metabolites and myokines, such as β-aminoisobutyric acid (BAIBA) and irisin, that enhance fuel oxidation in adipose tissue ^70–72^. In turn, BAT-derived myostatin has been reported to restrain skeletal muscle performance and exercise capacity as part of the homeostatic feedback axis ^73^. By analogy, cold induces hepatic production and release of acylcarnitines and triacylglycerol-rich VLDL particles that support FAO and thermogenesis in BAT ^32,74–76^. Reciprocally, the present study suggests that 3-OHSA is generated and released by BAT as a byproduct of active FAO and functions as a unique biological signal to the liver. Despite its lower circulating concentration than fatty acids, 3-OHSA potently alleviates ETC overload and reduces ROS production. In this model, BAT-derived 3-OHSA acts as a feedback mediator, protecting the liver from oxidative stress during high thermogenic demand. Recent studies have expanded the list of BAT-derived peptides that affect liver metabolism, including neuregulin 4, IL-6, and IL-1Ra, the secreted isoform of IL-1β antagonist ^13,77,78^. It has also been reported that BAT releases peptidase M20 domain-containing 1 (PM20D1), which generates N-acyl amino acids in circulation ^79^. N-acyl amino acids function as endogenous mitochondrial uncouplers, increasing respiration in various cell types, elevating whole-body energy expenditure, and potently reducing body weight in mice ^79^. In contrast, 3-OHSA primarily acts on the liver, reducing excess ROS production, in part through ANT2, and protecting the liver from oxidative stress. These studies support a BAT-liver axis that couples fuel provision with redox control in a mutually beneficial manner to control systemic metabolic health under conditions, such as cold exposure, where FAO demand is high across metabolic organs.

### Limitations of the study

First, the metabolite datasets analyzed here are limited to metabolites and lipids that could be reliably annotated using existing databases and available chemical standards. There are many unannotated features in the raw data that may represent key biomarkers of BAT activity and bioactive mediators derived from BAT. Future work should structurally characterize these metabolites and define their biological roles. Analyses of large sample sizes with a variety of metabolic conditions, such as obesity, diabetes, and aging, would also expand the list of BAT-derived factors and their metabolic regulation. Second, the mouse model used in this study (UCP1-Cre × PPARγ KO) efficiently ablated interscapular and perirenal BAT depots; however, this model did not ablate UCP1-negative beige adipocytes residing in inguinal WAT and other depots ^80,81^. These UCP1-negative adipocytes can influence systemic energy balance through UCP1-independent mechanisms ^82–84^, although their quantitative contribution to the circulating metabolite pool is likely minor relative to canonical BAT depots. Third, our human cohorts are primarily composed of healthy young men (average 23.2 years old) with a limited sample size. Future studies are warranted to investigate associations between BAT-derived metabolites and the prevalence of Type 2 diabetes and cardiovascular disease across diverse populations. Lastly, an important future study is to quantitatively determine the extent to which 3-OHSA and other metabolites mediate the protective effect of BAT against hepatic oxidative stress.

## RESOURCE AVAILABILITY

### Lead Contact

Further information and requests for resources and reagents should be directed to and will be fulfilled by the lead contact, Shingo Kajimura.

### Materials availability

Mouse strains generated in this study are available upon request from the lead contact.

### Data and code availability

- The mass spectrometry proteomics data have been deposited to Metabolomics Workbench (https://www.metabolomicsworkbench.org), with a project ID PR002639 and a project DOI 10.21228/M8J26T.
- Source data used to generate all graphs presented in this study are available in Data S1.
- This paper does not report original code.
- Any additional information required to reanalyze the data reported in this work paper is available from the Lead Contact upon request.

## ACKNOWLEDGMENTS

We are grateful to Dr. Yu-Hua Tseng for sharing the immortalized human brown preadipocytes. We also thank Hiroshi Nishida, Yong Chen, Tadashi Yamamuro, Christopher Auger, Jihoon Shin, Kazusa Miyachi, and Daqing Wang for their insightful discussions, valuable advice, or technical support. This work was supported by grants from the National Institutes of Health (NIH) (DK125283, DK097441, DK126160 to SK), Howard Hughes Medical Institute to SK, the Japan Agency for Medical Research and Development (JP20gm1310007 to TY, JS), and the Japan Science and Technology Agency (JPMJFR2014 to TY).

## AUTHOR CONTRIBUTIONS

Conceptualization, D.W., S.K.; methodology, D.W., M.L., T.L., M.M., M.S., T.Y.; investigation, D.W., M.L., T.L., M.M., T.Y.; formal analysis, D.W., T.Y.; resources, J.S., M.S., T.Y., S.K.; writing – draft writing, D.W., S.K.; review and editing, all the co-authors; funding acquisition, J.S., T.Y., and S.K.; supervision, J.S., M. S., T.Y., and S.K.

## STAR Method

### EXPERIMENTAL MODEL AND STUDY PARTICIPANT DETAILS

#### Animals

All animal experiments conducted were performed in compliance of protocols approved by the Institutional Animal Care and Use Committee at Beth Israel Deaconess Medical Center (028-2022). All mice were housed under a 12 h - 12 h light/dark cycle. Room-temperature mice were housed at 23 °C in ventilated cages with an ACH of 25. Mice housed at thermal neutral conditions were housed in an incubator at 28°C. Mice exposed to cold were individually housed in an incubator set to 6 °C under ad libitum conditions. Mice were fed a standard diet (Lab Diet 5008) or a high-fat diet (Research Diets; D12492; 60% Fat) and had free access to food and water, unless experimentally specified. Male C57BL/6J (Stock No. 000664) and male C57BL/6J DIO mice (Stock No. 380050) were purchased from the Jackson Laboratory. *PPARg*^flox/flox^ (Stock No. 004584), and *Ucp1*-Cre (Stock No. 024670) mice were obtained from the Jackson Laboratory. Mice supplemented with BSA-conjugated form of 3-hydroxy stearic acid (Cayman 22692) received a daily *i.p.* injection of 8 mg/kg/day. Control mice were supplemented with the equal amount of unconjugated BSA. 22 weeks DIO mice maintained on 60% Fat diet were used for *i.p.* injection experiment. 10-12 weeks male mice were used for all experiments unless otherwise specified.

To generate the BSA-conjugated form of the 3-hydroxy stearic acid, a 10% bovine serum albumin (BSA) solution was pre-incubated in a 37 °C water bath. The compound was dissolved at 250 mM in ethanol and heated to 65 °C. After 15 min, the concentrated compound solution was slowly added to the 10% BSA at a 1:100 dilution. The mixture was incubated at 37 °C for 2 h with gentle intermittent agitation until the solution turned from cloudy to clear. The final conjugate was sterilized by filtration through a 0.22 μm membrane and used for *i.p.* injection.

#### Human subjects

Healthy adult East Asian (Japanese) volunteers (mean age 23.2 years, *n* = 84) were provided with information about the study and gave their written informed consent for participation in the FDG-PET/CT examination. The FDG-PET/CT data obtained in our previous studies were enrolled in the analysis through an opt-out process ^85^. The FDG-PET/CT examinations were performed in the winter with a fasting for ∼12 h ^85^. The participants were exposed to a standardized no shivering cold exposure at 19°C for 1 hour, received ^18^F-FDG (1.7 MBq kg−1 body weight) intravenously and remained in the same cold conditions for another hour. Cold-induced BAT activity was quantified by SUV of FDG in the supraclavicular adipose deposits with Hounsfield Units from −300 to −10. The protocols were approved by the Institutional Research Ethics Review Board of Tenshi College in Sapporo, Japan (Protocol# 2015-25) and the University of Tokyo, Japan (Protocol# 23-82). Information on the subjects is shown in **Supplementary Table 2**.

To stratify high BAT vs. low BAT subjects, we used median-split dichotomization, which is considered an unbiased statistical method, particularly when no cut-off values are established, and this method has been widely used in ^18^FDG-PET-based BAT studies ^29,45–49^. Note that there is no clear consensus for a biologically meaningful cut-off value for BAT in the field. For example, SUV>2.0 has been proposed to distinguish active BAT from WAT or blood pool ^86^, while SUV>1.5 has been used as a cut-off value to compare BAT volume ^14,44^. Although we attempted to use cut-off values of 2.0 or 1.5, this led to significant skew in the sample size and an inequitable analysis. When we applied the SUV cut-off of >2, 58 participants (67%) were classified as BAT+, while only 28 (33%) were classified as BAT-. The skew becomes larger when we use the cut-off value SUV > 1.5. Accordingly, we used median-split dichotomization. The cut-off value of SUV≥3.4 is aligned with a previous study with SUV > 4.0 ^29^.

#### Cells

For mouse brown adipocyte culture, the base media was high glucose DMEM (Gibco 11965092), containing 10% FBS (R&D systems S11550) and 1% P/S (Gibco 15140). To induce differentiation in brown adipocytes, confluent preadipocytes were treated with an induction cocktail consisting of 0.5 mM isobutylmethylxanthine (Sigma I5879), 125 μM indomethacin (Sigma I7378), 2 μg/mL dexamethasone (Sigma D4902), 850 nM insulin (Sigma I6634), and 1 nM T3 (Sigma T2877) for 2 days. Following induction, the cells were switched to a maintenance medium of base medium supplemented with 850 nM insulin and 1 nM T3. The cells were kept in the maintenance medium for 4-6 days to achieve full differentiation.

For human brown adipocyte culture, preadipocytes were plated and grown in high glucose DMEM medium (Gibco 11965092) supplemented with 10% FBS and 1% P/S. For adipocyte differentiation, cells were grown for 3 days until reaching confluence, and then treated with the induction medium containing 2% FBS, 0.5 mM isobutylmethylxanthine, 0.1 μM dexamethasone, 0.5 μM human insulin, 2 nM T3, 30 μM indomethacin, 17 μM pantothenate, 33 μM biotin (Sigma-Aldrich, Dallas, TX) for another 12 days, with medium refreshed every 3 days.

### METHOD DETAILS

#### Mouse tissue and tissue explant

For tissue isolation, mice were euthanized and were perfused by PBS through the heart, target tissues were rapidly dissected, rinsed in ice-cold sterile PBS to remove residual blood, blotted dry, and immediately snap-frozen in liquid nitrogen. Samples were then stored at -80 °C until LC-MS study.

For tissue explant culture, mice were euthanized and were perfused with PBS through the heart, target tissues were rapidly excised, rinsed in ice-cold sterile PBS, and transferred to DMEM/F-12 (Gibco 11320033) supplemented with 10% FBS and 1% P/S. Tissues were minced into ∼1-2 mm³ pieces. The pieces were washed with sterile PBS. 0.1 g pieces were resuspended in 0.5 ml fresh medium (DMEM/F12 + 10% FBS + 1% P/S). Medium was replaced after 2 h to remove surgery-related debris. Finally, medium is collected at 0h, 12h, 24h, and 36h, followed by centrifugation at ∼500 × g for 10 min to remove cell debris and particulates.

#### Extracellular fluid isolation

Brown adipose tissue and epidydimal white adipose tissue were dissected from adult wild-type male mice (Jax 000664) and placed in the center of 20 μm nylon net filter (Millipore NY2004700), which was secured in a 1.5 mL tube^87,88^. BAT was then centrifuged at 800 × g for 10 minutes at 4°C. The extracellular fluid collected from the centrifugation was snap-froze in liquid nitrogen and stored in -80°C until metabolite extraction.

#### Serum collection

For human serum collection, after fasting for ∼12 h, serum was collected from subjects who rested at 27°C. Subjects were then exposed to standardized nonshivering cold exposure at 19°C, and serum was collected after 1 hours at 19°C. For mouse serum collection, the mouse blood was collected from the tail into tubes containing clotting activator (Starstedt Inc, 16.440.100). The samples were kept on ice and the serum was separated by centrifugation at 3,000 g for 10 min at 4 °C.

#### Metabolite extraction

For serum samples, 4 volumes of cold methanol were added to 1 volume of serum, 2 volumes of 80% methanol were supplemented to obtain a better protein precipitation. For tissue samples, metabolites were extracted by homogenizing the tissues with 80% methanol at a 40:1 volume to wet weight ratio. The mixture was vigorously vortexed for 30 s, incubated at -80 °C for 1 hour and then centrifuged at 16000 × g for 15 min at 4 °C. The supernatant was carefully collected and stored at -80 °C until LC-MS analysis. Metabolite extracts were used for both metabolomics and hydrophilic lipidomics.

#### Lipid extraction

The Bligh & Dyer method, with slight optimization, was employed to extract lipids from the samples. For serum, cell media or extracellular fluid, 1 volume of samples was mixed with 20 volumes of methanol: chloroform: water (2:1:0.8, v/v/v) and vortexed thoroughly to precipitate proteins and release lipids. To induce phase separation, additional chloroform and water were added to achieve a final ratio of methanol: chloroform: water of 2:2:1.8 (v/v/v).

For tissue samples, 0.1 g tissue was homogenized with 100 μL of chloroform and 200 μL of methanol. To induce phase separation, additional 100 μL of chloroform and 100 μL of water were added to achieve a final ratio of methanol: chloroform: water of 2:2:1.8 (v/v/v).

The final mixture of the extraction buffer for serum, cell media, extracellular fluid or tissue samples was vortexed and centrifuged at 10,000 × g for 10 min at 4 °C. The lower organic phase was carefully collected, dried under vacuum, and reconstituted in acetonitrile: isopropanol (1:1, v/v) for LC-MS analysis. During extraction, SPLASH™ LIPIDOMIX™ Mass Spec Standard was added to monitor variability. Samples with a coefficient of variation (CV) greater than 0.3 were discarded and re-prepared.

#### LC-MS metabolomics

All LC-MS data are acquired on a UHPLC system (Vanquish Horizon, Thermo Scientific) coupled to an orbitrap mass spectrometer (Exploris 240, Thermo Scientific). An ACQUITY UPLC BEH Amide column (1.7 μm × 2.1 mm × 100 mm) (Waters Corporation, Milford, MA, USA) kept at 25 °C was employed for chromatographic separation. Mobile phases A = 25 mM ammonium acetate and 25 mM ammonium hydroxide in 100% water, and B = 100% acetonitrile. The linear gradient eluted from 95% B (0.0-1 min), 95% B to 65% B (1-7.0 min), 65% B to 40% B (7.0-8.0 min), 40% B (8.0-9.0 min), 40% B to 95% B (9.0-9.1 min), then stayed at 95% B for 5.9 min. The flow rate was 0.4 mL/min. ESI source parameters were set as follows: spray voltage, 3500 V or -2800 V, in positive or negative modes, respectively; vaporizer temperature, 350 °C; sheath gas, 50 arb; aux gas, 10 arb; ion transfer tube temperature, 325 °C. The full scan was set as: orbitrap resolution, 12,0000; RF Lens (%), 70; maximum injection time, 100 ms; scan range, 70-800 Da. The ddMS2 scan was set as: orbitrap resolution, 30,000; maximum injection time, 54 ms; isolation width, 1.5 m/z; HCD collision energy (%), 30, 50,150; Dynamic exclusion mode was set as auto. The metabolites were annotated by matching their retention times and MS² spectra against our in-house library and searching the MS² spectra against the mzCloud database.

#### LC-MS lipidomics

All LC-MS data are acquired on a UHPLC system (Vanquish Horizon, Thermo Scientific) coupled to an orbitrap mass spectrometer (Exploris 240, Thermo Scientific). For hydrophobic lipids, an AQUITY UPLC BEH C18 column (1.7 μm × 2.1 mm × 100 mm) (Waters Corporation, Milford, MA, USA) kept at 50 °C was employed for chromatographic separation. Mobile phases A = acetonitrile: water (60:40), and B= isopropanol: acetonitrile: water (90:8:2). Both mobile phases contained 0.1% formic acid and 10 mM ammonium formate. A stepwise gradient elution was performed as follows: 0-1 min 20% B, 1-3 min 20-30% B, 3-4 min 30-45% B, 4-6 min 45-60% B, 6-8 min 60-65% B, 8-10 min 65-65% B, 10-17 min 65-98% B, 17-21 min 98-98% B, 21-21min 98%-20% B, 21-25min 20%-20% B. Flow rate was 0.3

mL/min and the injection volume was 3 μL for positive mode and 5 μL for negative mode. The full scan was set as: orbitrap resolution, 12,0000; RF Lens (%), 70; maximum injection time, 100 ms; scan range, 200-1700 Da. The ddMS2 scan was set as: orbitrap resolution, 30,000; maximum injection time, 54 ms; isolation width, 1.5 m/z; HCD collision energies (%), 20,30,40 for negative mode and 25,30 for positive mode. Scan range mode was set with the first mass defined at m/z 75. Lipids were identified and quantified by MSDIAL 5.5 and Lipidsearch software.

For hydrophilic lipids, an ACQUITY UPLC HSS T3 Column (100Å, 1.8 µm, 2.1 mm × 100 mm) (Waters Corporation, Milford, MA, USA) kept at 25 °C was employed for chromatographic separation. Mobile phase A was water with 0.1% formic acid and mobile phase B was acetonitrile with 0.1% formic acid. A stepwise gradient elution was performed as follows: 0-1 min 1% B, 1-8 min 1-100% B, 8-10 min 100-100% B, 10-10.1 min 100-1% B, 10.1-12 min 1-1% B. Flow rate was 0.4 mL/min. ESI source parameters were set as follows: spray voltage, 3500 V or -2800 V, in positive or negative modes, respectively; vaporizer temperature, 350 °C; sheath gas, 50 arb; aux gas, 10 arb; ion transfer tube temperature, 325 °C. The full scan was set as: orbitrap resolution, 12,0000; RF Lens (%), 70; maximum injection time, 100 ms; scan range, 70-800 Da. The ddMS2 scan was set as: orbitrap resolution, 30,000; maximum injection time, 54 ms; isolation width, 1.5 m/z; HCD collision energy (%), 30, 50,150; Dynamic exclusion mode was set as auto. Lipids were identified and quantified by MSDIAL 5.5 and Lipidsearch software.

#### BAT-linked circulating molecules screening

An unpaired *t*-test was used to determine the significance of each circulating metabolite or lipid between BAT-ablated mice and littermate controls under each temperature condition. Considering the correction methods such as Bonferroni or FDR in this setting would substantially increase the risk of Type II errors, potentially masking biologically relevant but modest effects, we used unadjusted p-values as an initial filter to generate a candidate list, rather than as definitive evidence of significance. A total of 206 metabolites and 249 lipids with p < 0.05 in at least one condition were selected as the candidates of BAT-responsive circulating molecules, then included for correlation clustering. Clustering analysis was performed using the Mfuzz package in R software, and pairwise Pearson correlation coefficients between molecules were used to generate heatmaps. **Table S1** lists the p-values and fold changes between the BAT-ablated mice and littermate controls for these metabolites and lipids.

#### Absolute concentrations measurement

The absolute concentrations of indicated fatty acids and 3-hydroxy stearic acid were determined using an external standard curve-based quantification approach. Standard curves spanning the physiological concentration ranges of the indicated molecules were generated. Samples and standards were analyzed within the same LC–MS batch. Concentrations of the indicated molecules in the injected samples were calculated using the linear regression equations derived from the corresponding standard curves. Final absolute concentrations in the original samples were determined by correcting for the dilution factors introduced during extraction.

#### Primary hepatocyte isolation

Briefly, mice were anesthetized, and the portal vein was cannulated for *in situ* liver perfusion. The liver was first perfused with pre-warmed Hanks’ balanced salt solution (HBSS) containing 16 mM HEPES, 25 mM NaHCO₃, 10 mM glucose, 0.5 mM EGTA, and 1% penicillin–streptomycin (P/S), pH 7.4. This was followed by perfusion with HBSS supplemented with 20 μg/mL Liberase TM, 16 mM HEPES, 25 mM NaHCO₃, 10 mM glucose, 5 mM CaCl₂·2H₂O, and 1% P/S, pH 7.4, until the liver parenchyma became soft. The liver was then excised, gently dissociated, and filtered to obtain a single-cell suspension. Hepatocytes were collected by centrifugation at 100 × g for 1 min at 4 °C, washed twice with ice-cold William’s E medium, and resuspended in William’s E medium supplemented with 10% FBS, 1% P/S, and 1% GlutaMAX. An equal volume of 70% Percoll was added, mixed gently, and centrifuged at 624 × g for 10 min at 4 °C. The resulting pellet was washed once and resuspended in William’s E medium containing 10% FBS, 1% P/S, and 1% GlutaMAX.

#### ANT2 RNAi in primary hepatocyte

Primary hepatocytes were isolated as described above and seeded to ∼80% confluence in William’s E medium containing 10% FBS, 1% P/S, and 1% GlutaMAX. Cells were transfected with siRNA targeting *Slc25a5* (ANT2) at a final concentration of 1 nM using Lipofectamine RNAiMAX, according to the manufacturer’s instructions. Briefly, siRNA and Lipofectamine RNAiMAX were separately diluted in Opti-MEM medium, combined at a 1:1 ratio, and incubated for 5 min at room temperature to allow complex formation. The siRNA–lipid complexes were then added directly to the cells. Cells were incubated for 48 h before downstream analyses. Knockdown efficiency of ANT2 was assessed by qPCR using primers listed in **Table S5**.

#### Primary hepatocyte respiration

Isolated primary hepatocytes were loaded into Oroboros O2k chambers at 0.5 × 10⁶ cells·mL⁻¹. Basal respiration was initiated with 10 mM glucose, 1 mM pyruvate and 2 mM glutamine. Sequential additions included 2.5 μM oligomycin, 1 μM carbonyl cyanide-p-trifluoromethoxyphenylhydrazone (FCCP), and 2 μM each of rotenone and antimycin A.

#### Mouse tissue lysate respiration

Mice were euthanized and were perfused with PBS through the heart, target tissues were rapidly dissected, rinsed in ice-cold sterile PBS to remove residual blood, and blotted dry. Approximately 0.05 g of each tissue was homogenized in 1 mL of mitochondrial isolation medium (MIM; 300 mM sucrose, 10 mM HEPES, 1 mM EGTA, pH 7.2). The lysates were then resuspended in buffer Z (100 mM MES potassium salt, 30 mM KCl, 10 mM KH₂PO₄, 5 mM MgCl₂, 1 mM EGTA) to a final concentration of ∼0.5 mg/mL in the Oroboros O2k chamber for respiration measurements. Respiration was initiated with 2.5 mM malate, 5 mM pyruvate, 10 mM glutamate, and 10 mM succinate, followed by stimulation with 5 μM 3-OHSA.

#### Mitochondrial isolation

Mice were anesthetized, and the liver was perfused in situ with ice-cold PBS via the portal vein to remove circulating blood. Liver was homogenized in mitochondrial isolation media (MIM) containing 300 mM sucrose, 10 mM HEPES, 1 mM EGTA, 1 mg/mL fatty acid-free BSA at pH 7.2. Homogenized samples were centrifuged at 800 × g for 5 min at 4°C, and the supernatant was transferred to a new tube and pelleted at 10000 × g for 10 min at 4°C. Pelleted mitochondria were resuspended in homogenization buffer without BSA and normalized for equal protein through the BCA assay.

#### Mitochondrial H₂O₂ production rate

H₂O₂ production rate was measured using a plate reader with excitation at 550 nm and emission at 590 nm at 37 °C in a final volume of 200 μL per well. A 2× H₂O₂ reaction buffer, containing 5 mM malate, 10 mM pyruvate, 20 mM glutamate, 20 mM succinate, 6 U/mL horseradish peroxidase (HRP), and 20 μM Amplex® Red, was prepared in buffer Z. For each well, 100 μL of the 2× H₂O₂ buffer was mixed with 100 μL of 2× mitochondrial suspension (∼0.2 mg/mL in buffer Z) immediately before plate scanning to initiate the reaction. *J*H₂O₂ was determined as the rate of change over a specified period.

#### Mitochondrial membrane potential

In Safranin O assays ^89^, fluorescence measurements were performed in 2 mL of standard reaction medium containing 250 mM sucrose, 10 mM HEPES, 200 µM EGTA, 2 mM NaH₂PO₄, 1 mM MgCl₂, 2.5 mM malate, 5 mM pyruvate, 10 mM glutamate, and 10 mM succinate, supplemented with 5 µM Safranine O. Measurements were conducted in a cuvette at 37 °C with continuous stirring. Fluorescence was recorded using a Fluoromax fluorometer (HORIBA) in kinetic mode with excitation at 495 nm and emission at 586 nm (bandwidth 5 nm). After establishing a baseline, mitochondria were added to a final concentration of 0.5 mg/mL. For the measurement of ΔΨm in primary hepatocytes, fluorescence reading was performed in standard reaction medium supplemented with 40 μM digitonin and 5 μM safranine O. Approximately 4 × 10⁶ cells were collected, centrifuged at 1,500 × g for 3 min, and washed once with standard reaction medium. Cell pellets were resuspended in 1 mL of standard reaction medium containing 40 μM digitonin and incubated for 5 min. Cells were then centrifuged (1,500 × g, 3 min) and resuspended in standard reaction medium to a final volume of 50 μL. After baseline stabilization, cells were added to the cuvette at a final concentration of 2 × 10⁶ cells/mL.

At the midpoint of the ΔΨm polarization range, 3-OHSA or other control lipids were added. The rate of decrease in fluorescence intensity (CPS per second) was calculated to assess the rate of ΔΨm polarization. For inhibitor experiments, ANT or mPTP inhibitors were added fifty seconds after the addition of 3-OHSA or other lipids to examine their effects on 3-OHSA- or lipids-induced reductions of ΔΨm polarization rate.

#### Mitochondrial respiration

Mitochondrial respiration was assessed using Oroboros O2k oxygraphs. Isolated mitochondria were suspended in buffer Z containing 100 mM MES potassium salt, 30 mM KCl, 10 mM KH₂PO₄, 5 mM MgCl₂, and 1 mM EGTA. The final liver mitochondrial concentration was ∼0.1 mg/mL. Basal respiration was initiated with 2.5 mM malate, 5 mM pyruvate, 10 mM glutamate, and 10 mM succinate. Sequential additions included 2.5 μM oligomycin, 60 nM carbonyl cyanide-p-trifluoromethoxyphenylhydrazone (FCCP), and 2 μM each of rotenone and antimycin A.

#### Mitochondrial ATP production rate

ATP production rate was measured using a plate reader with excitation at 340 nm and emission at 460 nm (5 nm excitation band) at 37 °C in a final volume of 200 μL per well. A 2× ATP reaction buffer, containing 5 mM malate, 10 mM pyruvate, 20 mM glutamate, 20 mM succinate, 2 U/mL hexokinase, 4 U/mL glucose-6-phosphate dehydrogenase, 4 mM glucose and 4 mM NADP, was prepared in buffer Z. For each well, 100 μL of the 2× ATP buffer was mixed with 100 μL of 2× mitochondrial suspension (∼0.2 mg/mL in buffer Z) immediately before plate scanning to initiate the reaction. *J*ATP was determined as the rate of change over a specified period.

#### Indirect calorimetry

22 weeks male DIO mice fed 60% HFD diet (D12492; 60% Fat) were singly housed and monitored with Promethion Metabolic Cage System (Sable Systems). Mice received a daily *i.p.* injection of BSA-conjugated form of 3-hydroxy stearic acid (Cayman 22692) at a dose of 8 mg/kg/day. Control mice were supplemented with the equal amount of unconjugated BSA. Mice were initially housed at room temperature (23 °C) and then changed into thermoneutrality (30 °C). Mice had unrestricted access to food and water throughout the experiment. The average light and dark cycle measurements were calculated by taking the last full light or dark cycle prior to the temperature being changed. Data were analyzed with CalR (https://calrapp.org/).

#### Glucose and insulin tolerance

22 weeks male DIO mice fed 60% HFD diet were housed at room temperature and received a daily *i.p.* injection of BSA-conjugated form of 3-hydroxy stearic acid (Cayman 22692) at a dose of 8 mg/kg/day for 2 weeks. Unconjugated BSA and BSA-conjugated stearic acid at the same concentrations were used as controls. Mice were fasted for 4 hours prior to intraperitoneal delivery of glucose (1g per kg body mass) or insulin (1 U per kg body mass). Blood glucose from the tail vein was measured repeatedly from 0 to 120 minutes post-injection with a handheld glucometer (Abbott, Freestyle Lite).

#### Oxidative stress markers

Protein oxidative stress accumulation in the liver was determined through protein carbonyl content, which was measured using a commercially available kit (Abcam, ab126287). Assay was performed with 30 mg of tissue, and the counts were normalized to protein content, which was determined by a BCA assay. Lipid oxidative stress in the liver was determined by measuring malondialdehyde (MDA) with a commercially available kit (Abcam, ab118970) utilizing colormetric assay and 10 mg of tissue. For the oxidative stress marker assessment in BAT-ablation mice, a cohort of mice aged approximately 60 weeks was used.

#### Quantitative RT-PCR (qPCR)

Total RNA was isolated from cells or tissue using Trizol (Invitrogen) according to manufacturer instructions. RNA was reverse transcribed using iScript cDNA synthesis kit (Biorad). PCR reactions were performed with Applied Biosystems QuantStudio 6 Flex using Sybrgreen (Biorad). Assays were performed in four replicates, and all results were normalized to β-actin, which was unchanged between controls and respective experimental groups. All values are relative to the mean of the control group. Mitochondrial DNA copy number was quantified by qPCR using mitochondrial genes 16S (F: CCGCAAGGGAAAGATGAAAGAC; R: TCGTTTGGTTTCGGGGTTTC) and ND1 (F: CTAGCAGAAACAAACCGGGC; R: CCGGCTGCGTATTCTACGTT) and normalized to the nuclear gene HK2 (F: GCCAGCCTCTCCTGATTTTAGTGT; R: GGGAACACAAAAGACCTCTTCTGG). All primers used are listed in **Supplementary Table 5**.

## QUANTIFICATION AND STATISTICAL ANALYSIS

### Statistics

Unless otherwise indicated, Data were expressed as mean ± SD. Statistical analysis was performed using GraphPad Prism 8 (GraphPad Software, Inc., La Jolla, Ca). *P* values below 0.05 were considered statistically significant throughout the study. When two-group comparisons were performed, a two-sample unpaired Student’s *t*-test was used unless otherwise indicated. For Figure 4H, two-way ANOVA with Šídák’s correct for multiple comparisons was used.

**Supplemental table 1. P value, fold change, and cluster result of serum metabolites and lipids altered in BAT-ablated mice model. M, metabolites cluster; L, lipids cluster. Fold change, BAT+/BAT-. (Related to Figure 1)**

**Supplemental table 2. Clinical parameters of human participants. (Related to Figure 3)**

**Supplemental table 3. Cold-induced lipid changes of human subjects (*p* < 0.05) (Related to Figure 3)**

**Supplemental table 4. Difference of cold-induced lipid changes between high and low BAT groups (*p* < 0.05) (Related to Figure 3)**

**Supplemental Table 5. Primer sequences used for quantitative RT-PCR. (Related to Figure S1, S5 and S6)**

## REFERENCES

1. Cannon, B., and Nedergaard, J. (2004). Brown adipose tissue: function and physiological significance. Physiological reviews 84, 277–359. 10.1152/physrev.00015.2003.

2. Betz, M.J., and Enerback, S. (2018). Targeting thermogenesis in brown fat and muscle to treat obesity and metabolic disease. Nat Rev Endocrinol 14, 77–87. 10.1038/nrendo.2017.132.

3. Koenen, M., Hill, M.A., Cohen, P., and Sowers, J.R. (2021). Obesity, Adipose Tissue and Vascular Dysfunction. Circ Res 128, 951–968. 10.1161/CIRCRESAHA.121.318093.

4. Sakers, A., De Siqueira, M.K., Seale, P., and Villanueva, C.J. (2022). Adipose-tissue plasticity in health and disease. Cell 185, 419–446. 10.1016/j.cell.2021.12.016.

5. Carpentier, A.C., Blondin, D.P., Haman, F., and Richard, D. (2023). Brown Adipose Tissue-A Translational Perspective. Endocr Rev 44, 143–192. 10.1210/endrev/bnac015.

6. Ohno, H., Shinoda, K., Ohyama, K., Sharp, L.Z., and Kajimura, S. (2013). EHMT1 controls brown adipose cell fate and thermogenesis through the PRDM16 complex. Nature 504, 163–167. 10.1038/nature12652.

7. Cohen, P., Levy, J.D., Zhang, Y., Frontini, A., Kolodin, D.P., Svensson, K.J., Lo, J.C., Zeng, X., Ye, L., Khandekar, M.J., et al. (2014). Ablation of PRDM16 and beige adipose causes metabolic dysfunction and a subcutaneous to visceral fat switch. Cell 156, 304–316. 10.1016/j.cell.2013.12.021.

8. Hasegawa, Y., Ikeda, K., Chen, Y., Alba, D.L., Stifler, D., Shinoda, K., Hosono, T., Maretich, P., Yang, Y., Ishigaki, Y., et al. (2018). Repression of Adipose Tissue Fibrosis through a PRDM16-GTF2IRD1 Complex Improves Systemic Glucose Homeostasis. Cell metabolism 27, 180–194 e186. 10.1016/j.cmet.2017.12.005.

9. Enerback, S., Jacobsson, A., Simpson, E.M., Guerra, C., Yamashita, H., Harper, M.E., and Kozak, L.P. (1997). Mice lacking mitochondrial uncoupling protein are cold-sensitive but not obese. Nature 387, 90–94. 10.1038/387090a0.

10. Feldmann, H.M., Golozoubova, V., Cannon, B., and Nedergaard, J. (2009). UCP1 ablation induces obesity and abolishes diet-induced thermogenesis in mice exempt from thermal stress by living at thermoneutrality. Cell metabolism 9, 203–209. 10.1016/j.cmet.2008.12.014.

11. Rahbani, J.F., Bunk, J., Lagarde, D., Samborska, B., Roesler, A., Xiao, H., Shaw, A., Kaiser, Z., Braun, J.L., Geromella, M.S., et al. (2024). Parallel control of cold-triggered adipocyte thermogenesis by UCP1 and CKB. Cell metabolism 36, 526–540 e527. 10.1016/j.cmet.2024.01.001.

12. Verkerke, A.R.P., Wang, D., Yoshida, N., Taxin, Z.H., Shi, X., Zheng, S., Li, Y., Auger, C., Oikawa, S., Yook, J.S., et al. (2024). BCAA-nitrogen flux in brown fat controls metabolic health independent of thermogenesis. Cell 187, 2359–2374 e2318. 10.1016/j.cell.2024.03.030.

13. Wang, H., Kelly, M., Aziz, N., Tsagkaraki, E., Nicoloro, S.M., Lifshitz, L.M., Guilherme, A., Rowland, L.A., Yenilmez, B., Santos, K.B., et al. (2025). Thermogenic adipocytes alleviate hepatic steatosis and insulin resistance via macrophage cytokine secretion in obese mice. BioRxiv. 10.1101/2025.08.01.667482.

14. Becher, T., Palanisamy, S., Kramer, D.J., Eljalby, M., Marx, S.J., Wibmer, A.G., Butler, S.D., Jiang, C.S., Vaughan, R., Schoder, H., et al. (2021). Brown adipose tissue is associated with cardiometabolic health. Nature medicine 27, 58–65. 10.1038/s41591-020-1126-7.

15. Raiko, J., Orava, J., Savisto, N., and Virtanen, K.A. (2020). High Brown Fat Activity Correlates With Cardiovascular Risk Factor Levels Cross-Sectionally and Subclinical Atherosclerosis at 5-Year Follow-Up. Arterioscler Thromb Vasc Biol 40, 1289–1295. 10.1161/ATVBAHA.119.313806.

16. Herz, C.T., Kulterer, O.C., Prager, M., Schmoltzer, C., Langer, F.B., Prager, G., Marculescu, R., Kautzky-Willer, A., Hacker, M., Haug, A.R., and Kiefer, F.W. (2021). Active Brown Adipose Tissue is Associated With a Healthier Metabolic Phenotype in Obesity. Diabetes. 10.2337/db21-0475.

17. Kajimura, S., Spiegelman, B.M., and Seale, P. (2015). Brown and Beige Fat: Physiological Roles beyond Heat Generation. Cell metabolism 22, 546–559. 10.1016/j.cmet.2015.09.007.

18. Cohen, P., and Kajimura, S. (2021). The cellular and functional complexity of thermogenic fat. Nature reviews. Molecular cell biology 22, 393–409. 10.1038/s41580-021-00350-0.

19. Villarroya, F., Cereijo, R., Villarroya, J., and Giralt, M. (2017). Brown adipose tissue as a secretory organ. Nat Rev Endocrinol 13, 26–35. 10.1038/nrendo.2016.136.

20. Scheele, C., and Wolfrum, C. (2020). Brown Adipose Crosstalk in Tissue Plasticity and Human Metabolism. Endocr Rev 41, 53–65. 10.1210/endrev/bnz007.

21. Yang, F.T., and Stanford, K.I. (2022). Batokines: Mediators of Inter-Tissue Communication (a Mini-Review). Curr Obes Rep 11, 1–9. 10.1007/s13679-021-00465-7.

22. Hondares, E., Iglesias, R., Giralt, A., Gonzalez, F.J., Giralt, M., Mampel, T., and Villarroya, F. (2011). Thermogenic activation induces FGF21 expression and release in brown adipose tissue. The Journal of biological chemistry 286, 12983–12990. 10.1074/jbc.M110.215889.

23. Ameka, M., Markan, K.R., Morgan, D.A., BonDurant, L.D., Idiga, S.O., Naber, M.C., Zhu, Z., Zingman, L.V., Grobe, J.L., Rahmouni, K., and Potthoff, M.J. (2019). Liver Derived FGF21 Maintains Core Body Temperature During Acute Cold Exposure. Sci Rep 9, 630. 10.1038/s41598-018-37198-y.

24. Lynes, M.D., Leiria, L.O., Lundh, M., Bartelt, A., Shamsi, F., Huang, T.L., Takahashi, H., Hirshman, M.F., Schlein, C., Lee, A., et al. (2017). The cold-induced lipokine 12,13-diHOME promotes fatty acid transport into brown adipose tissue. Nature medicine 23, 631–637. 10.1038/nm.4297.

25. Stanford, K.I., Lynes, M.D., Takahashi, H., Baer, L.A., Arts, P.J., May, F.J., Lehnig, A.C., Middelbeek, R.J.W., Richard, J.J., So, K., et al. (2018). 12,13-diHOME: An Exercise-Induced Lipokine that Increases Skeletal Muscle Fatty Acid Uptake. Cell metabolism 27, 1111–1120 e1113. 10.1016/j.cmet.2018.03.020.

26. Deshmukh, A.S., Peijs, L., Beaudry, J.L., Jespersen, N.Z., Nielsen, C.H., Ma, T., Brunner, A.D., Larsen, T.J., Bayarri-Olmos, R., Prabhakar, B.S., et al. (2019). Proteomics-Based Comparative Mapping of the Secretomes of Human Brown and White Adipocytes Reveals EPDR1 as a Novel Batokine. Cell metabolism 30, 963–975 e967. 10.1016/j.cmet.2019.10.001.

27. Cataldo, L.R., Gao, Q., Argemi-Muntadas, L., Hodek, O., Cowan, E., Hladkou, S., Gheibi, S., Spegel, P., Prasad, R.B., Eliasson, L., et al. (2022). The human batokine EPDR1 regulates beta-cell metabolism and function. Mol Metab 66, 101629. 10.1016/j.molmet.2022.101629.

28. Cypess, A.M., Cannon, B., Nedergaard, J., Kazak, L., Chang, D.C., Krakoff, J., Tseng, Y.H., Scheele, C., Boucher, J., Petrovic, N., et al. (2025). Emerging debates and resolutions in brown adipose tissue research. Cell metabolism 37, 12–33. 10.1016/j.cmet.2024.11.002.

29. Yoneshiro, T., Wang, Q., Tajima, K., Matsushita, M., Maki, H., Igarashi, K., Dai, Z., White, P.J., McGarrah, R.W., Ilkayeva, O.R., et al. (2019). BCAA catabolism in brown fat controls energy homeostasis through SLC25A44. Nature 572, 614–619. 10.1038/s41586-019-1503-x.

30. Duerre, D.J., Hansen, J.K., John, S.V., Jen, A., Carrillo, N.D., Bui, H., Bao, Y., Fabregat, M., Catrow, J.L., Chen, L.Y., et al. (2025). Haem biosynthesis regulates BCAA catabolism and thermogenesis in brown adipose tissue. Nat Metab 7, 1018–1033. 10.1038/s42255-025-01253-6.

31. Yoneshiro, T., Kataoka, N., Walejko, J.M., Ikeda, K., Brown, Z., Yoneshiro, M., Crown, S.B., Osawa, T., Sakai, J., McGarrah, R.W., et al. (2021). Metabolic flexibility via mitochondrial BCAA carrier SLC25A44 is required for optimal fever. eLife 10. 10.7554/eLife.66865.

32. Bartelt, A., Bruns, O.T., Reimer, R., Hohenberg, H., Ittrich, H., Peldschus, K., Kaul, M.G., Tromsdorf, U.I., Weller, H., Waurisch, C., et al. (2011). Brown adipose tissue activity controls triglyceride clearance. Nature medicine 17, 200–205. 10.1038/nm.2297.

33. Berbee, J.F., Boon, M.R., Khedoe, P.P., Bartelt, A., Schlein, C., Worthmann, A., Kooijman, S., Hoeke, G., Mol, I.M., John, C., et al. (2015). Brown fat activation reduces hypercholesterolaemia and protects from atherosclerosis development. Nature communications 6, 6356. 10.1038/ncomms7356.

34. Oh, C.M., Namkung, J., Go, Y., Shong, K.E., Kim, K., Kim, H., Park, B.Y., Lee, H.W., Jeon, Y.H., Song, J., et al. (2015). Regulation of systemic energy homeostasis by serotonin in adipose tissues. Nature communications 6, 6794. 10.1038/ncomms7794.

35. Suchacki, K.J., Ramage, L.E., Kwok, T.C., Kelman, A., McNeill, B.T., Rodney, S., Keegan, M., Gray, C., MacNaught, G., Patel, D., et al. (2023). The serotonin transporter sustains human brown adipose tissue thermogenesis. Nat Metab 5, 1319–1336. 10.1038/s42255-023-00839-2.

36. Crane, J.D., Palanivel, R., Mottillo, E.P., Bujak, A.L., Wang, H., Ford, R.J., Collins, A., Blumer, R.M., Fullerton, M.D., Yabut, J.M., et al. (2015). Inhibiting peripheral serotonin synthesis reduces obesity and metabolic dysfunction by promoting brown adipose tissue thermogenesis. Nat Med 21, 166–172. 10.1038/nm.3766.

37. Bornstein, M.R., Neinast, M.D., Zeng, X., Chu, Q., Axsom, J., Thorsheim, C., Li, K., Blair, M.C., Rabinowitz, J.D., and Arany, Z. (2023). Comprehensive quantification of metabolic flux during acute cold stress in mice. Cell metabolism 35, 2077–2092 e2076. 10.1016/j.cmet.2023.09.002.

38. Straat, M.E., Martinez-Tellez, B., Sardjoe Mishre, A., Verkleij, M.M.A., Kemmeren, M., Pelsma, I.C.M., Alcantara, J.M.A., Mendez-Gutierrez, A., Kooijman, S., Boon, M.R., and Rensen, P.C.N. (2022). Cold-Induced Thermogenesis Shows a Diurnal Variation That Unfolds Differently in Males and Females. The Journal of clinical endocrinology and metabolism 107, 1626–1635. 10.1210/clinem/dgac094.

39. Yau, W.W., Wong, K.A., Zhou, J., Thimmukonda, N.K., Wu, Y., Bay, B.H., Singh, B.K., and Yen, P.M. (2021). Chronic cold exposure induces autophagy to promote fatty acid oxidation, mitochondrial turnover, and thermogenesis in brown adipose tissue. iScience 24, 102434. 10.1016/j.isci.2021.102434.

40. Park, G., Haley, J.A., Le, J., Jung, S.M., Fitzgibbons, T.P., Korobkina, E.D., Li, H., Fluharty, S.M., Chen, Q., Spinelli, J.B., et al. (2023). Quantitative analysis of metabolic fluxes in brown fat and skeletal muscle during thermogenesis. Nat Metab 5, 1204–1220. 10.1038/s42255-023-00825-8.

41. Asantewaa, G., Tuttle, E.T., Ward, N.P., Kang, Y.P., Kim, Y., Kavanagh, M.E., Girnius, N., Chen, Y., Rodriguez, K., Hecht, F., et al. (2024). Glutathione synthesis in the mouse liver supports lipid abundance through NRF2 repression. Nat Commun 15, 6152. 10.1038/s41467-024-50454-2.

42. Yang, Y., Dieter, M.Z., Chen, Y., Shertzer, H.G., Nebert, D.W., and Dalton, T.P. (2002). Initial characterization of the glutamate-cysteine ligase modifier subunit Gclm(-/-) knockout mouse. Novel model system for a severely compromised oxidative stress response. J Biol Chem 277, 49446–49452. 10.1074/jbc.M209372200.

43. Chen, Y., Yang, Y., Miller, M.L., Shen, D., Shertzer, H.G., Stringer, K.F., Wang, B., Schneider, S.N., Nebert, D.W., and Dalton, T.P. (2007). Hepatocyte-specific Gclc deletion leads to rapid onset of steatosis with mitochondrial injury and liver failure. Hepatology 45, 1118–1128. 10.1002/hep.21635.

44. Chen, K.Y., Cypess, A.M., Laughlin, M.R., Haft, C.R., Hu, H.H., Bredella, M.A., Enerback, S., Kinahan, P.E., Lichtenbelt, W., Lin, F.I., et al. (2016). Brown Adipose Reporting Criteria in Imaging STudies (BARCIST 1.0): Recommendations for Standardized FDG-PET/CT Experiments in Humans. Cell metabolism 24, 210–222. 10.1016/j.cmet.2016.07.014.

45. Cameron, N.A., Petito, L.C., Colangelo, L.A., Gunderson, E.P., Catov, J.M., Grobman, W.A., Rana, J.S., Terry, J.G., Lloyd-Jones, D.M., Allen, N.B., and Khan, S.S. (2025). Prepregnancy Cardiovascular Health, Gestational Diabetes, and Coronary Artery Calcium. JAMA Cardiol 10, 888–895. 10.1001/jamacardio.2025.1887.

46. Zhang, X., Hack, L.M., Bertrand, C., Hilton, R., Gray, N.J., Boyar, L., Laudie, J., Heifets, B.D., Suppes, T., van Roessel, P.J., et al. (2025). Negative Affect Circuit Subtypes and Neural, Behavioral, and Affective Responses to MDMA: A Randomized Clinical Trial. JAMA Netw Open 8, e257803. 10.1001/jamanetworkopen.2025.7803.

47. Grkovski, M., Kohutek, Z.A., Schoder, H., Brennan, C.W., Tabar, V.S., Gutin, P.H., Zhang, Z., Young, R.J., Beattie, B.J., Zanzonico, P.B., et al. (2020). (18)F-Fluorocholine PET uptake correlates with pathologic evidence of recurrent tumor after stereotactic radiosurgery for brain metastases. Eur J Nucl Med Mol Imaging 47, 1446–1457. 10.1007/s00259-019-04628-6.

48. Unterrainer, M., Taugner, J., Kasmann, L., Tufman, A., Reinmuth, N., Li, M., Mittlmeier, L.M., Bartenstein, P., Kunz, W.G., Ricke, J., et al. (2022). Differential role of residual metabolic tumor volume in inoperable stage III NSCLC after chemoradiotherapy +/- immune checkpoint inhibition. Eur J Nucl Med Mol Imaging 49, 1407–1416. 10.1007/s00259-021-05584-w.

49. Carlier, T., Frecon, G., Mateus, D., Rizkallah, M., Kraeber-Bodere, F., Kanoun, S., Blanc-Durand, P., Itti, E., Le Gouill, S., Casasnovas, R.O., et al. (2024). Prognostic Value of (18)F-FDG PET Radiomics Features at Baseline in PET-Guided Consolidation Strategy in Diffuse Large B-Cell Lymphoma: A Machine-Learning Analysis from the GAINED Study. J Nucl Med 65, 156–162. 10.2967/jnumed.123.265872.

50. Li, J., Xu, J., Zhang, R., Hao, Y., He, J., Chen, Y., Jiao, G., and Abliz, Z. (2020). Strategy for Global Profiling and Identification of 2- and 3-Hydroxy Fatty Acids in Plasma by UPLC-MS/MS. Analytical chemistry 92, 5143–5151. 10.1021/acs.analchem.9b05627.

51. Wu, C., Orozco, C., Boyer, J., Leglise, M., Goodale, J., Batalov, S., Hodge, C.L., Haase, J., Janes, J., Huss, J.W., 3rd, and Su, A.I. (2009). BioGPS: an extensible and customizable portal for querying and organizing gene annotation resources. Genome biology 10, R130. 10.1186/gb-2009-10-11-r130.

52. Vyssokikh, M.Y., Holtze, S., Averina, O.A., Lyamzaev, K.G., Panteleeva, A.A., Marey, M.V., Zinovkin, R.A., Severin, F.F., Skulachev, M.V., Fasel, N., et al. (2020). Mild depolarization of the inner mitochondrial membrane is a crucial component of an anti-aging program. Proc Natl Acad Sci U S A 117, 6491–6501. 10.1073/pnas.1916414117.

53. Toime, L.J., and Brand, M.D. (2010). Uncoupling protein-3 lowers reactive oxygen species production in isolated mitochondria. Free Radic Biol Med 49, 606–611. 10.1016/j.freeradbiomed.2010.05.010.

54. Cadenas, S. (2018). Mitochondrial uncoupling, ROS generation and cardioprotection. Biochim Biophys Acta Bioenerg 1859, 940–950. 10.1016/j.bbabio.2018.05.019.

55. Suski, J., Lebiedzinska, M., Bonora, M., Pinton, P., Duszynski, J., and Wieckowski, M.R. (2018). Relation Between Mitochondrial Membrane Potential and ROS Formation. Methods Mol Biol 1782, 357–381. 10.1007/978-1-4939-7831-1_22.

56. Brookes, P.S. (2005). Mitochondrial H(+) leak and ROS generation: an odd couple. Free Radic Biol Med 38, 12–23. 10.1016/j.freeradbiomed.2004.10.016.

57. Mailloux, R.J., and Harper, M.E. (2012). Mitochondrial proticity and ROS signaling: lessons from the uncoupling proteins. Trends Endocrinol Metab 23, 451–458. 10.1016/j.tem.2012.04.004.

58. Bertholet, A.M., Chouchani, E.T., Kazak, L., Angelin, A., Fedorenko, A., Long, J.Z., Vidoni, S., Garrity, R., Cho, J., Terada, N., et al. (2019). H(+) transport is an integral function of the mitochondrial ADP/ATP carrier. Nature 571, 515–520. 10.1038/s41586-019-1400-3.

59. Bonora, M., Giorgi, C., and Pinton, P. (2022). Molecular mechanisms and consequences of mitochondrial permeability transition. Nat Rev Mol Cell Biol 23, 266–285. 10.1038/s41580-021-00433-y.

60. Niatsetskaya, Z., Sosunov, S., Stepanova, A., Goldman, J., Galkin, A., Neginskaya, M., Pavlov, E., and Ten, V. (2020). Cyclophilin D-dependent oligodendrocyte mitochondrial ion leak contributes to neonatal white matter injury. J Clin Invest 130, 5536–5550. 10.1172/JCI133082.

61. Alavian, K.N., Beutner, G., Lazrove, E., Sacchetti, S., Park, H.A., Licznerski, P., Li, H., Nabili, P., Hockensmith, K., Graham, M., et al. (2014). An uncoupling channel within the c-subunit ring of the F1FO ATP synthase is the mitochondrial permeability transition pore. Proc Natl Acad Sci U S A 111, 10580–10585. 10.1073/pnas.1401591111.

62. Harm, T., Dittrich, K., Brun, A., Fu, X., Frey, M., Petersen Uribe, A., Schwarz, F.J., Rohlfing, A.K., Castor, T., Geisler, T., et al. (2023). Large-scale lipidomics profiling reveals characteristic lipid signatures associated with an increased cardiovascular risk. Clin Res Cardiol 112, 1664–1678. 10.1007/s00392-023-02260-x.

63. Rivas Serna, I.M., Sitina, M., Stokin, G.B., Medina-Inojosa, J.R., Lopez-Jimenez, F., Gonzalez-Rivas, J.P., and Vinciguerra, M. (2021). Lipidomic Profiling Identifies Signatures of Poor Cardiovascular Health. Metabolites 11. 10.3390/metabo11110747.

64. Naderi, J., Johnson, A.K., Thakkar, H., Chandravanshi, B., Ksiazek, A., Anand, A., Vincent, V., Tran, A., Kalimireddy, A., Singh, P., et al. (2025). Ceramide-induced FGF13 impairs systemic metabolic health. Cell Metab 37, 1206–1222 e1208. 10.1016/j.cmet.2025.03.002.

65. Lopez, M., Dieguez, C., Tena-Sempere, M., and Gonzalez-Garcia, I. (2025). Ceramides in the central control of metabolism. Trends Endocrinol Metab 36, 11–14. 10.1016/j.tem.2024.06.007.

66. Summers, S.A., Chaurasia, B., and Holland, W.L. (2019). Metabolic Messengers: ceramides. Nat Metab 1, 1051–1058. 10.1038/s42255-019-0134-8.

67. Bikman, B.T., and Summers, S.A. (2011). Ceramides as modulators of cellular and whole-body metabolism. J Clin Invest 121, 4222–4230. 10.1172/JCI57144.

68. Wang, D.D., Toledo, E., Hruby, A., Rosner, B.A., Willett, W.C., Sun, Q., Razquin, C., Zheng, Y., Ruiz-Canela, M., Guasch-Ferre, M., et al. (2017). Plasma Ceramides, Mediterranean Diet, and Incident Cardiovascular Disease in the PREDIMED Trial (Prevencion con Dieta Mediterranea). Circulation 135, 2028–2040. 10.1161/CIRCULATIONAHA.116.024261.

69. Mantovani, A., Bonapace, S., Lunardi, G., Salgarello, M., Dugo, C., Canali, G., Byrne, C.D., Gori, S., Barbieri, E., and Targher, G. (2018). Association between plasma ceramides and inducible myocardial ischemia in patients with established or suspected coronary artery disease undergoing myocardial perfusion scintigraphy. Metabolism 85, 305–312. 10.1016/j.metabol.2018.05.006.

70. Bostrom, P., Wu, J., Jedrychowski, M.P., Korde, A., Ye, L., Lo, J.C., Rasbach, K.A., Bostrom, E.A., Choi, J.H., Long, J.Z., et al. (2012). A PGC1-alpha-dependent myokine that drives brown-fat-like development of white fat and thermogenesis. Nature 481, 463–468. 10.1038/nature10777.

71. Lee, P., Linderman, J.D., Smith, S., Brychta, R.J., Wang, J., Idelson, C., Perron, R.M., Werner, C.D., Phan, G.Q., Kammula, U.S., et al. (2014). Irisin and FGF21 are cold-induced endocrine activators of brown fat function in humans. Cell metabolism 19, 302–309. 10.1016/j.cmet.2013.12.017.

72. Roberts, L.D., Bostrom, P., O’Sullivan, J.F., Schinzel, R.T., Lewis, G.D., Dejam, A., Lee, Y.K., Palma, M.J., Calhoun, S., Georgiadi, A., et al. (2014). beta-Aminoisobutyric acid induces browning of white fat and hepatic beta-oxidation and is inversely correlated with cardiometabolic risk factors. Cell metabolism 19, 96–108. 10.1016/j.cmet.2013.12.003.

73. Kong, X., Yao, T., Zhou, P., Kazak, L., Tenen, D., Lyubetskaya, A., Dawes, B.A., Tsai, L., Kahn, B.B., Spiegelman, B.M., et al. (2018). Brown Adipose Tissue Controls Skeletal Muscle Function via the Secretion of Myostatin. Cell metabolism 28, 631–643 e633. 10.1016/j.cmet.2018.07.004.

74. Simcox, J., Geoghegan, G., Maschek, J.A., Bensard, C.L., Pasquali, M., Miao, R., Lee, S., Jiang, L., Huck, I., Kershaw, E.E., et al. (2017). Global Analysis of Plasma Lipids Identifies Liver-Derived Acylcarnitines as a Fuel Source for Brown Fat Thermogenesis. Cell metabolism 26, 509–522 e506. 10.1016/j.cmet.2017.08.006.

75. Hoeke, G., Kooijman, S., Boon, M.R., Rensen, P.C., and Berbee, J.F. (2016). Role of Brown Fat in Lipoprotein Metabolism and Atherosclerosis. Circ Res 118, 173–182. 10.1161/CIRCRESAHA.115.306647.

76. Auger, C., Nishida, H., Yuan, B., Silva, G.M., Fujimoto, M., Li, M., Katoh, D., Wang, D., Granath-Panelo, M., Shin, J., et al. (2026). Mitochondrial control of fuel switching via carnitine biosynthesis. Science (New York, N.Y 391, eady5532. 10.1126/science.ady5532.

77. Wang, G.X., Zhao, X.Y., Meng, Z.X., Kern, M., Dietrich, A., Chen, Z., Cozacov, Z., Zhou, D., Okunade, A.L., Su, X., et al. (2014). The brown fat-enriched secreted factor Nrg4 preserves metabolic homeostasis through attenuation of hepatic lipogenesis. Nature medicine 20, 1436–1443. 10.1038/nm.3713.

78. Qing, H., Desrouleaux, R., Israni-Winger, K., Mineur, Y.S., Fogelman, N., Zhang, C., Rashed, S., Palm, N.W., Sinha, R., Picciotto, M.R., et al. (2020). Origin and Function of Stress-Induced IL-6 in Murine Models. Cell 182, 372–387 e314. 10.1016/j.cell.2020.05.054.

79. Long, J.Z., Svensson, K.J., Bateman, L.A., Lin, H., Kamenecka, T., Lokurkar, I.A., Lou, J., Rao, R.R., Chang, M.R., Jedrychowski, M.P., et al. (2016). The Secreted Enzyme PM20D1 Regulates Lipidated Amino Acid Uncouplers of Mitochondria. Cell 166, 424–435. 10.1016/j.cell.2016.05.071.

80. Lasar, D., Rosenwald, M., Kiehlmann, E., Balaz, M., Tall, B., Opitz, L., Lidell, M.E., Zamboni, N., Krznar, P., Sun, W., et al. (2018). Peroxisome Proliferator Activated Receptor Gamma Controls Mature Brown Adipocyte Inducibility through Glycerol Kinase. Cell reports 22, 760–773. 10.1016/j.celrep.2017.12.067.

81. Puigserver, P., Wu, Z., Park, C.W., Graves, R., Wright, M., and Spiegelman, B.M. (1998). A cold-inducible coactivator of nuclear receptors linked to adaptive thermogenesis. Cell 92, 829–839. 10.1016/s0092-8674(00)81410-5.

82. Ikeda, K., Kang, Q., Yoneshiro, T., Camporez, J.P., Maki, H., Homma, M., Shinoda, K., Chen, Y., Lu, X., Maretich, P., et al. (2017). UCP1-independent signaling involving SERCA2b-mediated calcium cycling regulates beige fat thermogenesis and systemic glucose homeostasis. Nature medicine 23, 1454–1465. 10.1038/nm.4429.

83. Wang, T., Sharma, A.K., Wu, C., Maushart, C.I., Ghosh, A., Yang, W., Stefanicka, P., Kovanicova, Z., Ukropec, J., Zhang, J., et al. (2024). Single-nucleus transcriptomics identifies separate classes of UCP1 and futile cycle adipocytes. Cell metabolism. 10.1016/j.cmet.2024.07.005.

84. Vargas-Castillo, A., Sun, Y., Smythers, A.L., Grauvogel, L., Dumesic, P.A., Emont, M.P., Tsai, L.T., Rosen, E.D., Zammit, N.W., Shaffer, S.M., et al. (2024). Development of a functional beige fat cell line uncovers independent subclasses of cells expressing UCP1 and the futile creatine cycle. Cell metabolism. 10.1016/j.cmet.2024.07.002.

85. Yoneshiro, T., Matsushita, M., Fuse-Hamaoka, S., Kuroiwa, M., Kurosawa, Y., Yamada, Y., Arai, M., Wei, Y., Iida, M., Kuma, K., et al. (2025). Pre-fertilization-origin preservation of brown fat-mediated energy expenditure in humans. Nat Metab 7, 778–791. 10.1038/s42255-025-01249-2.

86. Yoneshiro, T., Aita, S., Matsushita, M., Kayahara, T., Kameya, T., Kawai, Y., Iwanaga, T., and Saito, M. (2013). Recruited brown adipose tissue as an antiobesity agent in humans. The Journal of clinical investigation 123, 3404–3408. 10.1172/JCI67803.

87. Mittenbuhler, M.J., Jedrychowski, M.P., Van Vranken, J.G., Sprenger, H.G., Wilensky, S., Dumesic, P.A., Sun, Y., Tartaglia, A., Bogoslavski, D., A, M., et al. (2023). Isolation of extracellular fluids reveals novel secreted bioactive proteins from muscle and fat tissues. Cell metabolism 35, 535–549 e537. 10.1016/j.cmet.2022.12.014.

88. Mills, E.L., Harmon, C., Jedrychowski, M.P., Xiao, H., Garrity, R., Tran, N.V., Bradshaw, G.A., Fu, A., Szpyt, J., Reddy, A., et al. (2021). UCP1 governs liver extracellular succinate and inflammatory pathogenesis. Nat Metab 3, 604–617. 10.1038/s42255-021-00389-5.

89. Figueira, T.R., Melo, D.R., Vercesi, A.E., and Castilho, R.F. (2012). Safranine as a fluorescent probe for the evaluation of mitochondrial membrane potential in isolated organelles and permeabilized cells. Methods Mol Biol 810, 103–117. 10.1007/978-1-61779-382-0_7.

