## Supplemental figures and legends for "Brown fat protects against hepatic oxidative stress by remodeling the circulating metabolome"

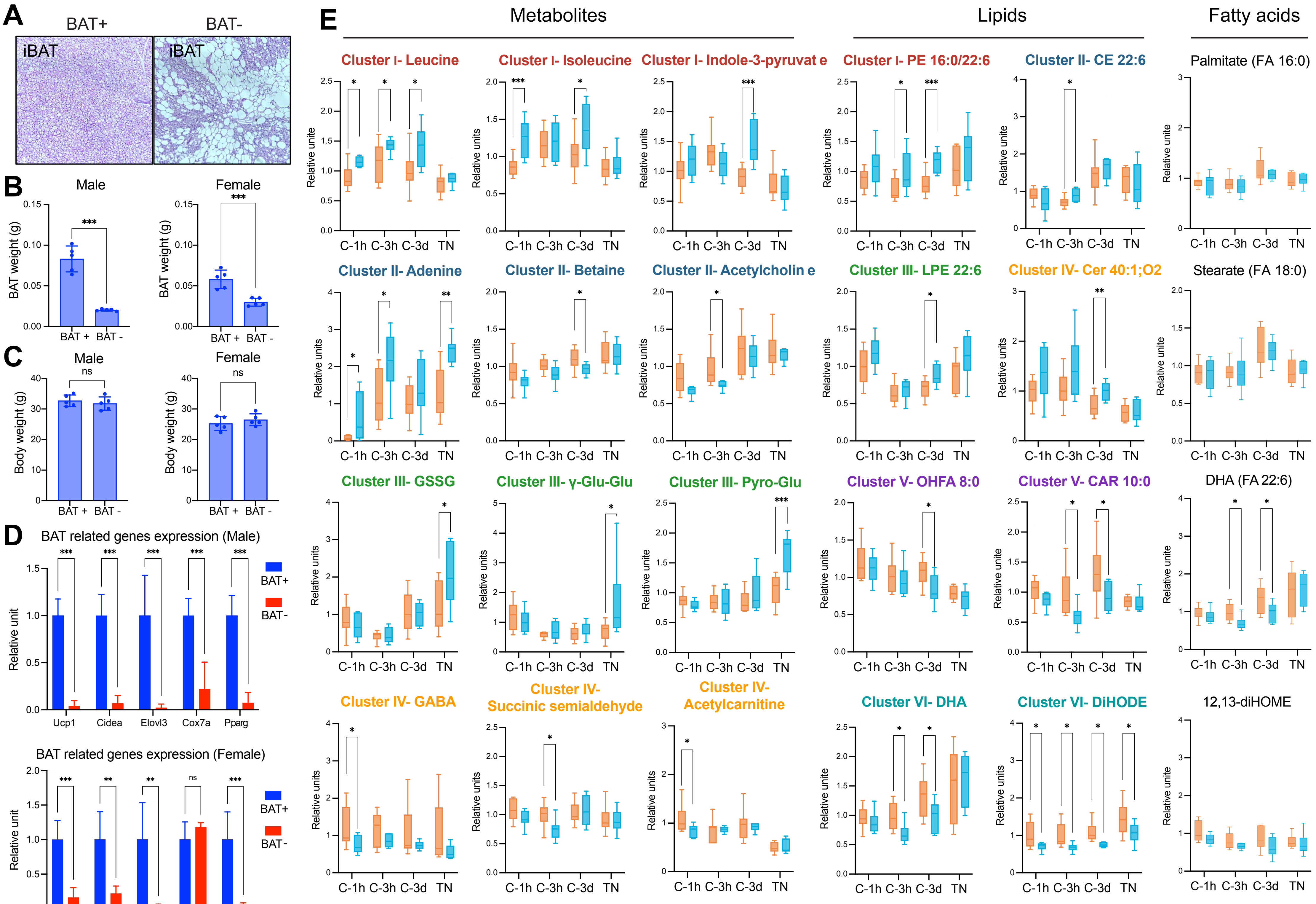
A
BAT+
BAT-
iBAT
iBAT
B
Male
Female
BAT weight (g)
BAT + BAT -
***
C
Male
Female
ns
Body weight (g)
BAT + BAT -
D
BAT related genes expression (Male)
BAT related genes expression (Female)
Relative unit
Ucp1
Cidea
Elovl3
Cox7a
Pparg
BAT+
BAT-
E
Metabolites
Lipids
Fatty acids
Cluster I- Leucine
Cluster I- Isoleucine
Cluster I- Indole-3-pyruvat e
Cluster I- PE 16:0/22:6
Cluster II- CE 22:6
Palmitate (FA 16:0)
Cluster II- Adenine
Cluster II- Betaine
Cluster II- Acetylcholin e
Cluster III- LPE 22:6
Cluster IV- Cer 40:1;O2
Stearate (FA 18:0)
Cluster III- GSSG
Cluster III- γ-Glu-Glu
Cluster III- Pyro-Glu
Cluster V- OHFA 8:0
Cluster V- CAR 10:0
DHA (FA 22:6)
Cluster IV- GABA
Cluster IV- Succinic semialdehyde
Cluster IV- Acetylcarnitine
Cluster VI- DHA
Cluster VI- DiHODE
12,13-diHOME
Relative units
Relative unite
C-1h C-3h C-3d TN
BAT+
BAT-

Figure S1

**Supplementary Figure 1. The impact of BAT on circulating metabolome and lipididome *in vivo* (Related to Figure 1)**

**A.** Morphology and hematoxylin and eosin (H&E) staining of interscapular BAT of BAT-ablated mice (BAT-) and littermate controls (BAT+).

**B-C.** Interscapular BAT weight (**B**) and body weight (**C**) measured in male (left) and female (right) BAT-ablated mice and their littermate controls. $n = 5$ in each group. Data are shown as mean ± SD. Statistic: unpaired *t*-test, $^{***}p < 0.001$.

**D.** Relative mRNA expression of BAT-related genes in BAT from male (up) and female (down) BAT-ablated mice and their littermate controls, measured by qPCR. $n = 5$ in each group. Data are shown as mean ± SD. Statistic: unpaired *t*-test, $^{*}p < 0.05$; $^{**}p < 0.01$; $^{***}p < 0.001$.

**E.** Quantitative profiles of the metabolites and lipids related to Figure 1. $n = 8$ for BAT-ablated mice and $n = 10$ for littermate controls. Statistic: unpaired *t*-test, $^{*}p < 0.05$; $^{**}p < 0.01$; $^{***}p < 0.001$. C-1h, 6 °C for 1 hour; C-3h, 6 °C for 3 hours; C-3d, 6 °C for 3 days; TN-3d, 28 °C for 3 days.

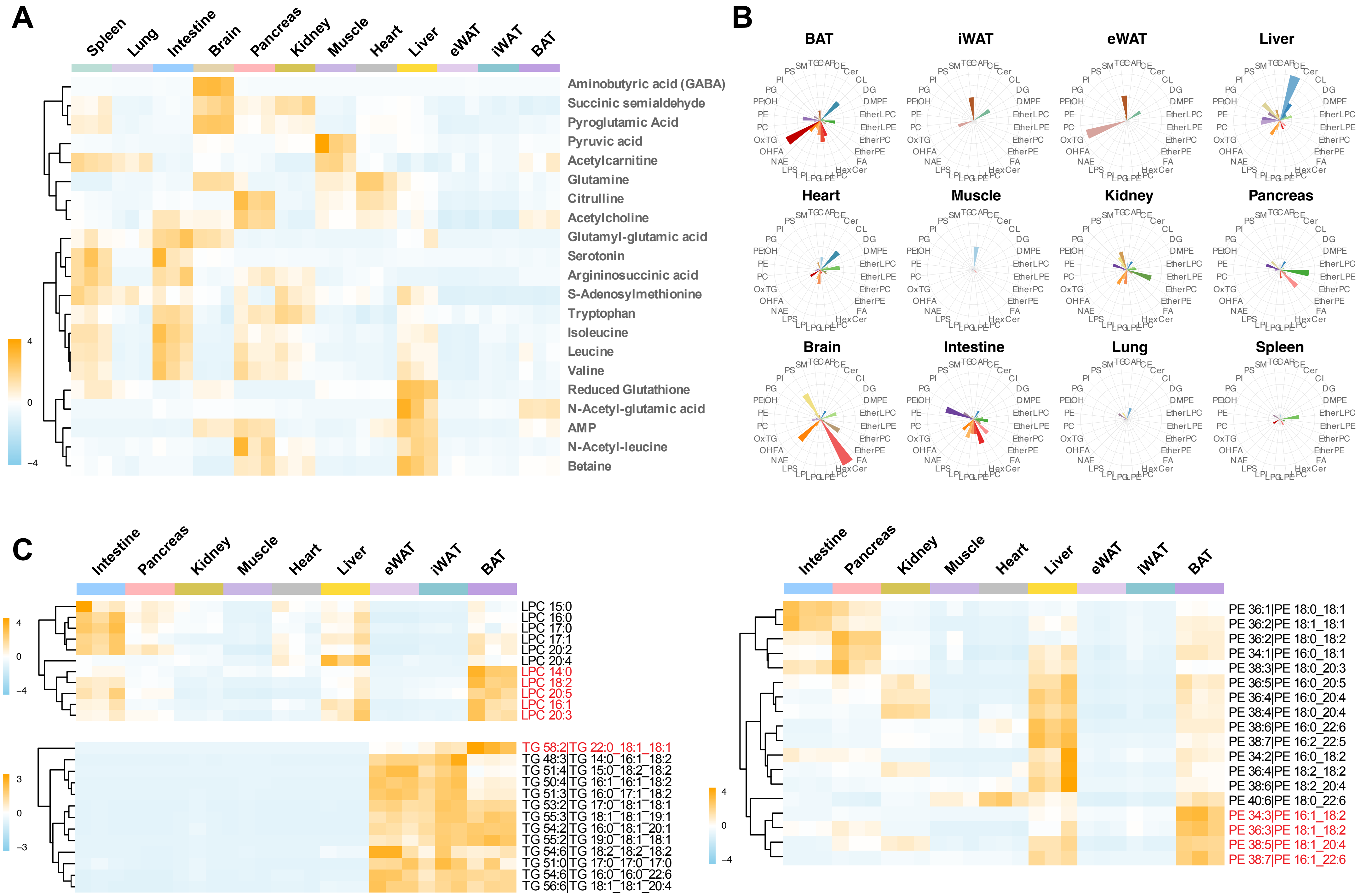
A
Spleen
Lung
Intestine
Brain
Pancreas
Kidney
Muscle
Heart
Liver
eWAT
iWAT
BAT
4
0
−4
Aminobutyric acid (GABA)
Succinic semialdehyde
Pyroglutamic Acid
Pyruvic acid
Acetylcarnitine
Glutamine
Citrulline
Acetylcholine
Glutamyl-glutamic acid
Serotonin
Argininosuccinic acid
S-Adenosylmethionine
Tryptophan
Isoleucine
Leucine
Valine
Reduced Glutathione
N-Acetyl-glutamic acid
AMP
N-Acetyl-leucine
Betaine
B
BAT
iWAT
eWAT
Liver
Heart
Muscle
Kidney
Pancreas
Brain
Intestine
Lung
Spleen
PS
SM
TG
CAR
CE
Cer
CL
DG
DMPE
EtherLPC
EtherLPE
EtherPC
EtherPE
FA
HexCer
PC
LPE
LPG
LPI
LPS
NAE
OHFA
OxTG
PC
PE
PEtOH
PG
PI
C
Intestine
Pancreas
Kidney
Muscle
Heart
Liver
eWAT
iWAT
BAT
4
0
−4
LPC 15:0
LPC 16:0
LPC 17:0
LPC 17:1
LPC 20:2
LPC 20:4
LPC 14:0
LPC 18:2
LPC 20:5
LPC 16:1
LPC 20:3
3
0
−3
TG 58:2|TG 22:0_18:1_18:1
TG 48:3|TG 14:0_16:1_18:2
TG 51:4|TG 15:0_18:2_18:2
TG 50:4|TG 16:1_16:1_18:2
TG 51:3|TG 16:0_17:1_18:2
TG 53:2|TG 17:0_18:1_18:1
TG 55:3|TG 18:1_18:1_19:1
TG 54:2|TG 16:0_18:1_20:1
TG 55:2|TG 19:0_18:1_18:1
TG 54:6|TG 18:2_18:2_18:2
TG 51:0|TG 17:0_17:0_17:0
TG 54:6|TG 16:0_16:0_22:6
TG 56:6|TG 18:1_18:1_20:4
Intestine
Pancreas
Kidney
Muscle
Heart
Liver
eWAT
iWAT
BAT
4
0
−4
PE 36:1|PE 18:0_18:1
PE 36:2|PE 18:1_18:1
PE 36:2|PE 18:0_18:2
PE 34:1|PE 16:0_18:1
PE 38:3|PE 18:0_20:3
PE 36:5|PE 16:0_20:5
PE 36:4|PE 16:0_20:4
PE 38:4|PE 18:0_20:4
PE 38:6|PE 16:0_22:6
PE 38:7|PE 16:2_22:5
PE 34:2|PE 16:0_18:2
PE 36:4|PE 18:2_18:2
PE 38:6|PE 18:2_20:4
PE 40:6|PE 18:0_22:6
PE 34:3|PE 16:1_18:2
PE 36:3|PE 18:1_18:2
PE 38:5|PE 18:1_20:4
PE 38:7|PE 16:1_22:6

Figure S2

**Supplementary Figure 2. Tissue origins of the BAT-responsive circulating metabolites and lipids (Related to Figure 2)**

**A.** Heatmap showing levels of BAT-responsive circulating metabolites across 12 mouse tissues.

**B.** Lipid source mapping of all quantified lipids across 12 mouse tissues. For each lipid class, the percentage of lipid species enriched in a specific tissue was calculated relative to the total number of enriched lipid species across all tissues. These percentages were then used to visualize the tissue-specific enrichment pattern of each lipid class. Lipids enriched in each tissue were identified using an unpaired *t*-test ($p < 0.05$) with fold change > 1, comparing tissue of interest to all other tissues. $n = 3$ for each tissue.

**C.** Distribution of carbon chain length and unsaturation degree of BAT-responsive circulating LPC, TG and PE lipid species across BAT, iWAT, eWAT, liver, heart, muscle, kidney, pancreas, intestine. Data represented as Z-score heatmap for indicated lipids. Orange represents a greater lipid abundance, and sky blue represents lesser lipid abundance. Brain, lung, and spleen, which did not show strong lipid enrichment, are not shown here. $n = 3$ per group. OHFA, hydroxy fatty acids; LPE, lysophosphatidylethanolamine.

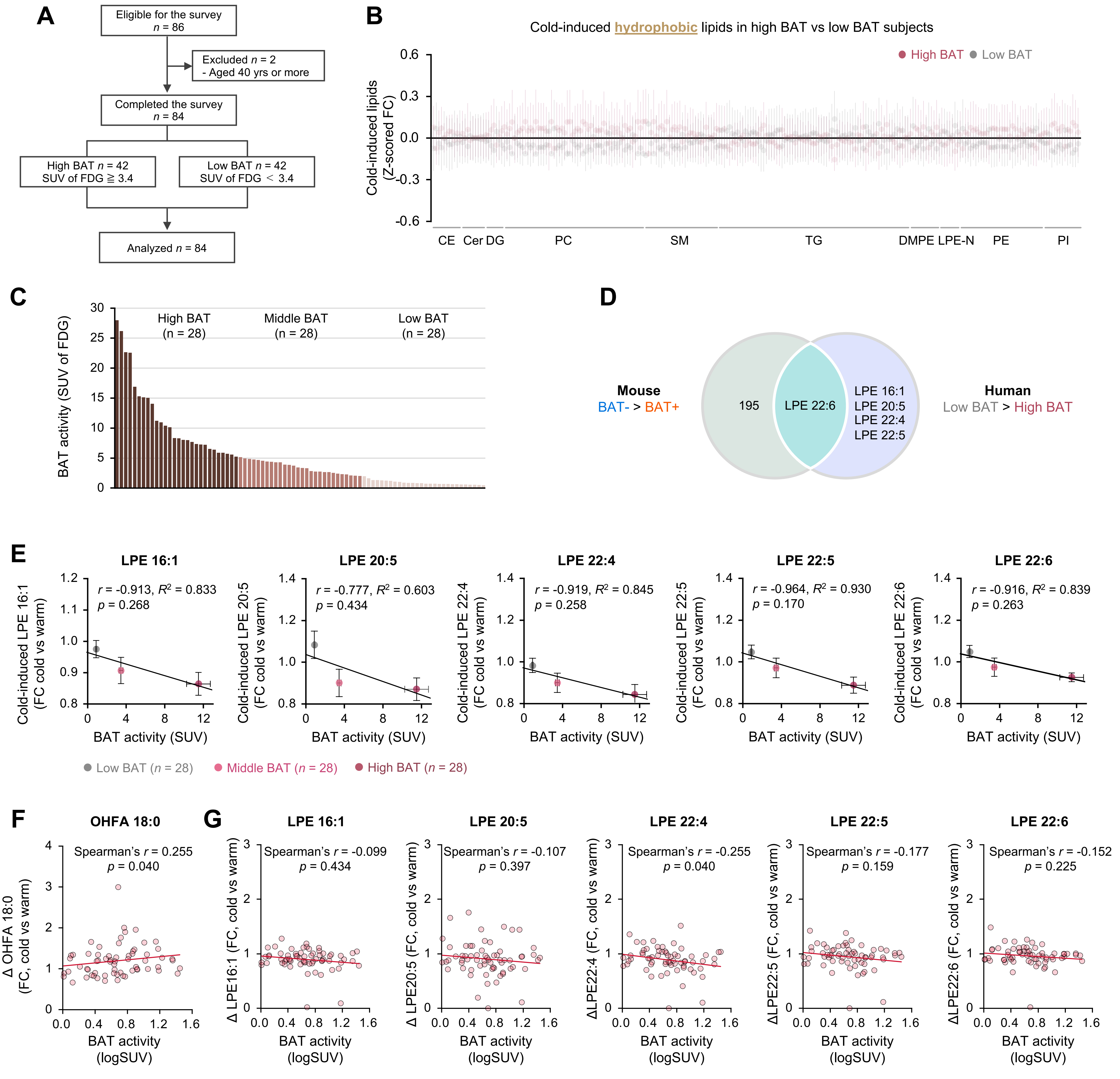

A
Eligible for the survey
n = 86
Excluded n = 2
- Aged 40 yrs or more
Completed the survey
n = 84
High BAT n = 42
SUV of FDG ≧ 3.4
Low BAT n = 42
SUV of FDG ＜ 3.4
Analyzed n = 84
B
Cold-induced hydrophobic lipids in high BAT vs low BAT subjects
High BAT
Low BAT
Cold-induced lipids (Z-scored FC)
CE
Cer
DG
PC
SM
TG
DMPE
LPE-N
PE
PI
C
BAT activity (SUV of FDG)
High BAT (n = 28)
Middle BAT (n = 28)
Low BAT (n = 28)
D
Mouse
BAT- > BAT+
195
LPE 22:6
LPE 16:1
LPE 20:5
LPE 22:4
LPE 22:5
Human
Low BAT > High BAT
E
LPE 16:1
r = -0.913, $R^2$ = 0.833
p = 0.268
Cold-induced LPE 16:1 (FC cold vs warm)
LPE 20:5
r = -0.777, $R^2$ = 0.603
p = 0.434
Cold-induced LPE 20:5 (FC cold vs warm)
LPE 22:4
r = -0.919, $R^2$ = 0.845
p = 0.258
Cold-induced LPE 22:4 (FC cold vs warm)
LPE 22:5
r = -0.964, $R^2$ = 0.930
p = 0.170
Cold-induced LPE 22:5 (FC cold vs warm)
LPE 22:6
r = -0.916, $R^2$ = 0.839
p = 0.263
Cold-induced LPE 22:6 (FC cold vs warm)
BAT activity (SUV)
Low BAT (n = 28)
Middle BAT (n = 28)
High BAT (n = 28)
F
OHFA 18:0
Spearman's r = 0.255
p = 0.040
Δ OHFA 18:0 (FC, cold vs warm)
G
LPE 16:1
Spearman's r = -0.099
p = 0.434
Δ LPE16:1 (FC, cold vs warm)
LPE 20:5
Spearman's r = -0.107
p = 0.397
Δ LPE20:5 (FC, cold vs warm)
LPE 22:4
Spearman's r = -0.255
p = 0.040
ΔLPE22:4 (FC, cold vs warm)
LPE 22:5
Spearman's r = -0.177
p = 0.159
ΔLPE22:5 (FC, cold vs warm)
LPE 22:6
Spearman's r = -0.152
p = 0.225
ΔLPE22:6 (FC, cold vs warm)
BAT activity (logSUV)


Figure S3

**Supplementary Figure 3. Cross-species integration of BAT-linked lipids in mice and humans (Related to Figure 3)**

**A.** CONSORT diagram of the human cohort. A total of 86 subjects were enrolled in this study. Two subjects aged 40 years old or more were excluded to minimize potential confounding effects of age. The final analysis included 84 healthy adults, who were divided into high- and low-BAT groups based on the median SUV value of 3.4 as the cutoff value. High BAT, SUV $\geqq$ 3.4, $n = 42$; low BAT, SUV < 3.4, $n = 42$.

**B.** Difference of cold-induced changes in hydrophobic lipids between the high BAT and low BAT groups. Cold-induced changes, Z-scored fold change (cold/warm). $n = 39$ for high BAT, $n = 41$ for low BAT (three and one samples for high and low BAT, respectively, were not sufficient for the measurement). Statistic: unpaired *t*-test. $^*p < 0.05$. CE, cholesterol ester; Cer, ceramide; DG, diglyceride; PC, phosphatidylcholine; SM, sphingomyelin; TG, triglyceride; DMPE, dimyristoylphosphatidylethanolamine; LPE-N, N-acyl-lysophosphatidylethanolamine; PE, phosphatidylethanolamine; PI, phosphatidylinositol.

**C.** Cold-induced FDG uptake into BAT (SUV) of human participants assessed by FDG-PET/CT ($n$ =84). Participants were classified into three groups based on tertiles with equal sample sizes: high BAT ($n = 28$, SUV $\geqq$ 5.2), middle BAT ($n = 28$, 2.0 $\leqq$ SUV < 5.2), and low BAT ($n = 28$, SUV < 2.0).

**D.** Overlap analysis of lipids accumulated in low BAT humans versus high BAT humans and those accumulated in BAT-ablated mice compared to control mice. LPE, lysophosphatidylethanolamine.

**E.** Correlation between FDG uptake in BAT and cold-induced fold change (FC) in the indicated circulating molecules. Statistic: correlation analysis and coefficient of determination ($R^2$). Participants were classified into three groups according to the tertile: high BAT ($n = 28$, SUV $\geqq$ 5.2), middle BAT ($n = 28$, 2.0 $\leqq$ SUV < 5.2), and low BAT groups ($n = 28$, SUV < 2.0).

**F.** Correlation between FDG uptake in BAT and cold-induced fold change (FC) of circulating OHFA 18:0 in subjects with SUV$\geqq$1 ($n = 65$). Spearman's rank correlation analysis was performed using individual participants.

**G.** Correlation between FDG uptake in BAT and cold-induced fold change (FC) of indicated circulating lipids in subjects with SUV$\geqq$1 ($n = 65$). Spearman's rank correlation analysis was performed using individual participants.

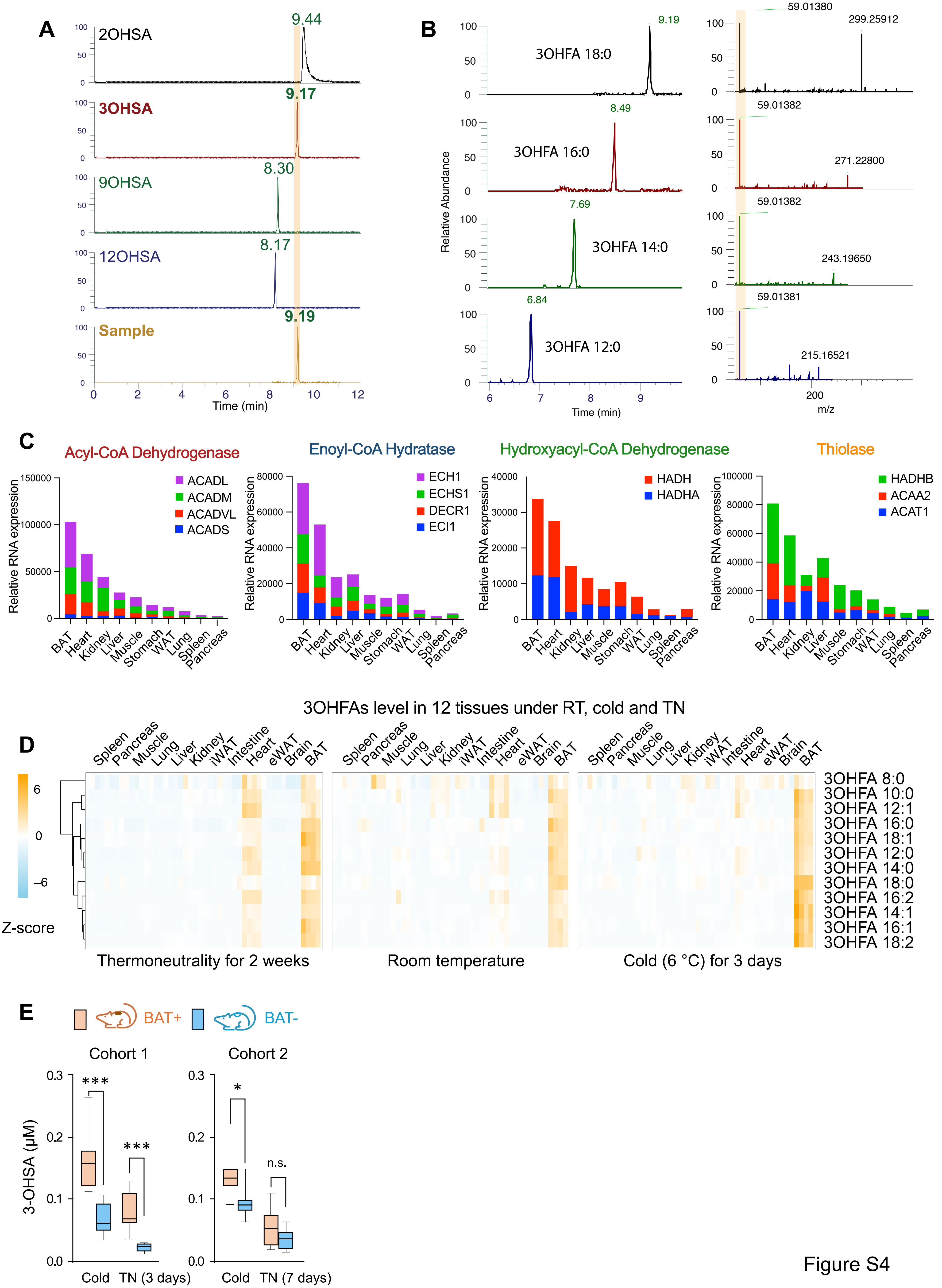

A
2OHSA
9.44
3OHSA
9.17
9OHSA
8.30
12OHSA
8.17
Sample
9.19
Time (min)
B
3OHFA 18:0
9.19
3OHFA 16:0
8.49
3OHFA 14:0
7.69
3OHFA 12:0
6.84
Relative Abundance
Time (min)
59.01380
299.25912
59.01382
271.22800
59.01382
243.19650
59.01381
215.16521
m/z
C
Acyl-CoA Dehydrogenase
ACADL
ACADM
ACADVL
ACADS
Enoyl-CoA Hydratase
ECH1
ECHS1
DECR1
ECI1
Hydroxyacyl-CoA Dehydrogenase
HADH
HADHA
Thiolase
HADHB
ACAA2
ACAT1
Relative RNA expression
BAT Heart Kidney Liver Muscle Stomach WAT Lung Spleen Pancreas
3OHFAs level in 12 tissues under RT, cold and TN
D
Spleen Pancreas Muscle Lung Liver Kidney iWAT Intestine Heart eWAT Brain BAT
Z-score
3OHFA 8:0
3OHFA 10:0
3OHFA 12:1
3OHFA 16:0
3OHFA 18:1
3OHFA 12:0
3OHFA 14:0
3OHFA 18:0
3OHFA 16:2
3OHFA 14:1
3OHFA 16:1
3OHFA 18:2
Thermoneutrality for 2 weeks
Room temperature
Cold (6 °C) for 3 days
E
BAT+
BAT-
Cohort 1
Cohort 2
3-OHSA (μM)
***
***
*
n.s.
Cold
TN (3 days)
TN (7 days)


Figure S4

**Supplementary Figure 4. 3-Hydroxy stearic acid (3-OHSA) is a cold-inducible BAT-derived lipid released into circulation**

**(Related to Figure 4)**

**A.** Determination of hydroxyl group positions in hydroxy stearic acid by comparing retention times with chemically synthesized standards (2-, 3-, 9-, and 12-hydroxy stearic acid).

**B.** Identification of hydroxyl group positions in other hydroxy fatty acids based on MS/MS diagnostic fragments (m/z 59.0138).

**C.** Relative RNA expression level of each fatty acid oxidation enzyme across 10 mouse tissues. Data was obtained from the BioGPS resource. Enzyme colors correspond to those in Figure 4B. Step 1, Acyl-CoA Dehydrogenase: ACADL, acyl-CoA dehydrogenase, long chain; ACADM, acyl-CoA dehydrogenase, medium chain; ACADVL, acyl-CoA dehydrogenase, very long chain; ACADS, acyl-CoA dehydrogenase, short chain; Step 2, Enoyl-CoA Hydratase: ECH1, enoyl-CoA hydratase 1; ECHS1, enoyl-CoA hydratase, short chain 1; DECR1, 2,4-dienoyl CoA reductase 1, for unsaturated fatty acids; ECI1, enoyl-CoA delta isomerase 1, for unsaturated fatty acids; Step3, Hydroxyacyl-CoA Dehydrogenase: HADH, hydroxyacyl-CoA dehydrogenase, medium to short chain; HADHA, hydroxyacyl-CoA dehydrogenase trifunctional multienzyme subunit α, long chain; Step 4, Thiolase: HADHB, hydroxyacyl-CoA dehydrogenase trifunctional multienzyme subunit β, long chain; ACAA2, acetyl-CoA acyltransferase 2, medium to long chain; ACAT1, Acetyl-CoA acetyltransferase 1, medium to long chain.

**D.** Heatmap of 3-hydroxy fatty acid levels across 12 mouse tissues under room temperature (RT), cold exposure (6 °C, 3 days), and thermoneutrality (TN, 28 °C, 2 weeks). Data represented as Z-score heatmap for indicated lipids. Orange represents a greater lipid abundance, and sky blue represents lesser lipid abundance. $n = 4$ per group.

**E.** Absolute concentrations of 3-OHSA in serum from two independent cohorts of BAT-ablated mice and littermate controls following 3 days of cold exposure or thermoneutrality. Left, cohort 1, $n = 8$ for BAT-ablated mice and $n = 10$ for controls; right, cohort 2, $n = 8$ per group. Mice were kept at room temperature and then exposed to cold (6 °C for 3 days), and subsequently acclimated to thermoneutrality (28 °C for 3 days for cohort 1 and 28 °C for 7 days for cohort 2). Statistic: unpaired *t*-test, $^{*}p < 0.05$; $^{**}p < 0.01$; $^{***}p < 0.001$.

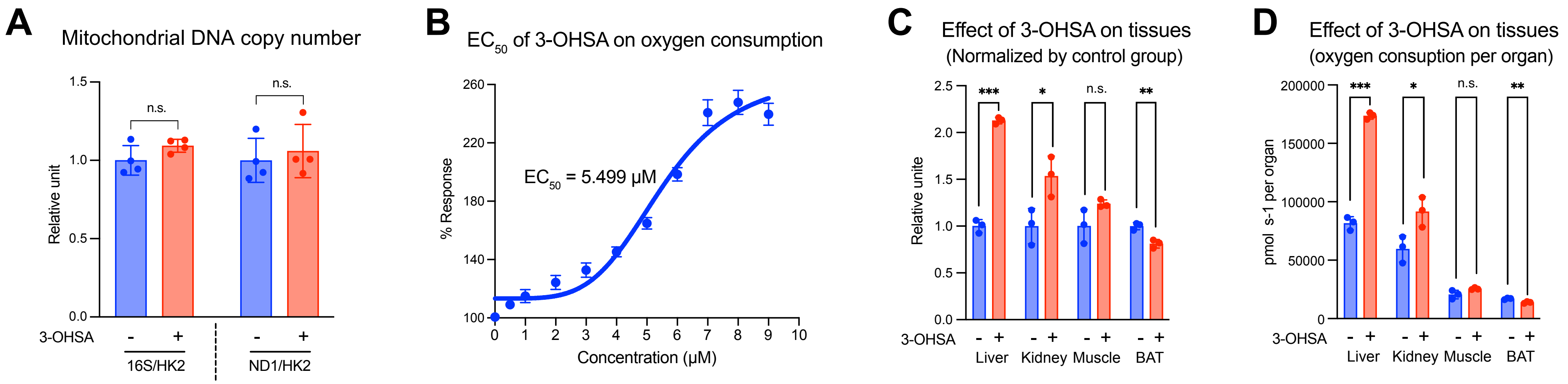


**Supplementary Figure 5. 3-hydroxy stearic acid causes mitochondrial inefficiency in the liver (Related to Figure 5)**

**A.** Mitochondrial DNA copy number was quantified by qPCR of mitochondrial genes (16S and ND1) and normalized to the nuclear gene HK2 under basal and 3-OHSA treatment conditions. Values are normalized to the basal condition. $n$ = 4. Data are shown as mean ± SD, with individual values presented. Statistic: unpaired *t*-test.

**B.** $EC_{50}$ of 3-hydroxy stearic acid on liver mitochondrial respiration. $EC_{50}$ value is determined by non-linear regression fitting of dose-response curves (3-OHSA vs. response, variable slope, four parameters). $n$ = 3.

**C-D.** Effect of 3-OHSA on tissue lysate respiration from the indicated metabolic organs. **C**, Respiration is presented as relative rates normalized to the vehicle-stimulated control group. **D,** Respiration is normalized to total organ mass to estimate whole-organ respiratory capacity under vehicle and 3-OHSA stimulation. $n$ = 3. Data are shown as mean ± SD.

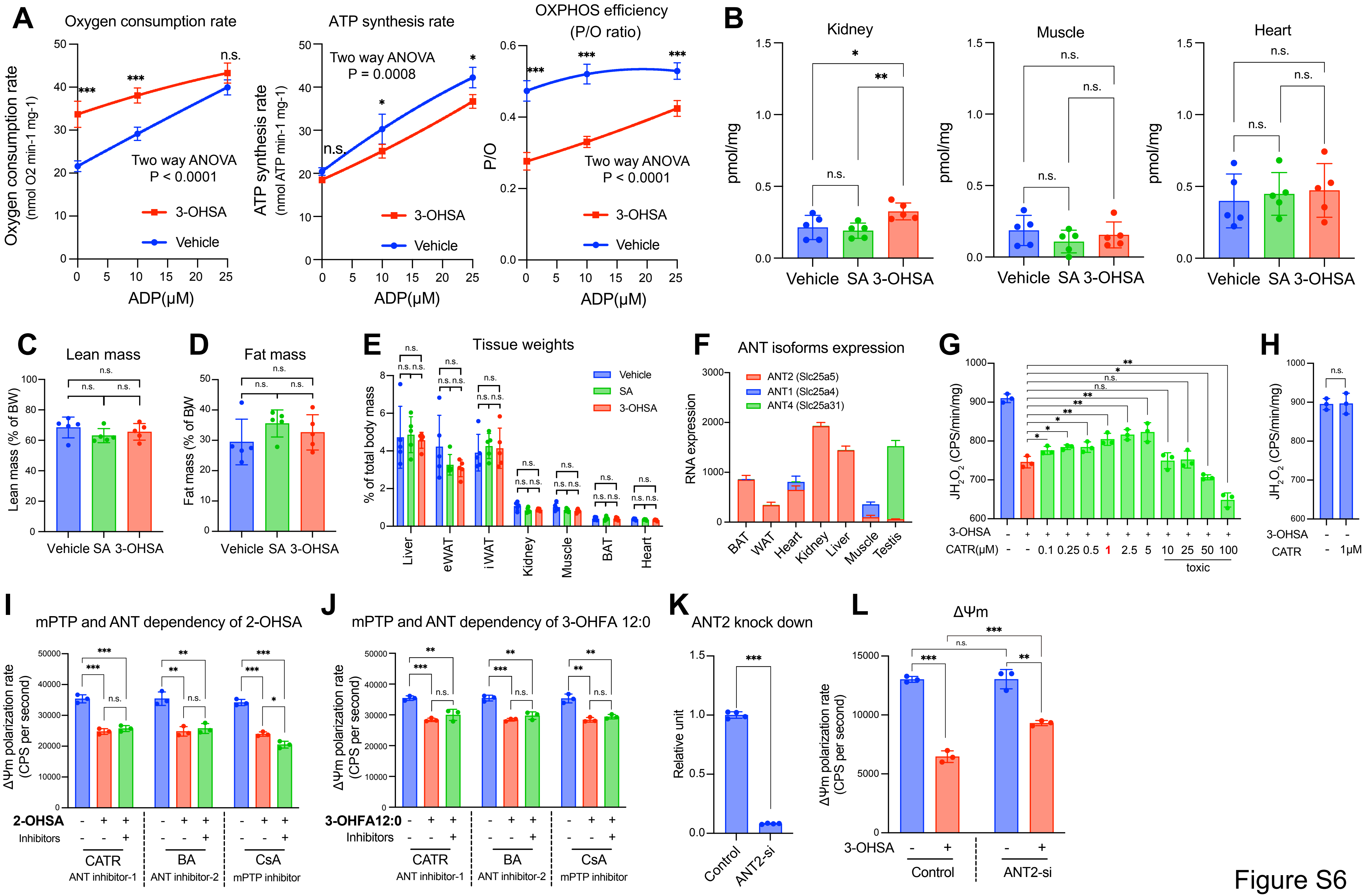

A
Oxygen consumption rate
Oxygen consumption rate (nmol O2 min-1 mg-1)
Two way ANOVA P < 0.0001
3-OHSA
Vehicle
ADP(μM)
ATP synthesis rate
ATP synthesis rate (nmol ATP min-1 mg-1)
Two way ANOVA P = 0.0008
OXPHOS efficiency (P/O ratio)
P/O
Two way ANOVA P < 0.0001
B
Kidney
Muscle
Heart
pmol/mg
Vehicle SA 3-OHSA
C
Lean mass
Lean mass (% of BW)
D
Fat mass
Fat mass (% of BW)
E
Tissue weights
% of total body mass
Liver eWAT iWAT Kidney Muscle BAT Heart
F
ANT isoforms expression
ANT2 (Slc25a5)
ANT1 (Slc25a4)
ANT4 (Slc25a31)
RNA expression
BAT WAT Heart Kidney Liver Muscle Testis
G
JH2O2 (CPS/min/mg)
3-OHSA
CATR(μM) 0.1 0.25 0.5 1 2.5 5 10 25 50 100
toxic
H
CATR 1μM
I
mPTP and ANT dependency of 2-OHSA
ΔΨm polarization rate (CPS per second)
2-OHSA
Inhibitors
CATR ANT inhibitor-1
BA ANT inhibitor-2
CsA mPTP inhibitor
J
mPTP and ANT dependency of 3-OHFA 12:0
3-OHFA12:0
K
ANT2 knock down
Relative unit
Control
ANT2-si
L
ΔΨm
Control
ANT2-si

Figure S6

**Supplementary Figure 6. 3-hydroxy stearic acid reduces hepatic mitochondrial membrane potential and oxidative stress.**

**(Related to Figure 6)**

**A.** Effect of 3-hydroxy stearic acid (3-OHSA) on oxygen consumption rate (left), ATP synthesis rate (middle), and OXPHOS efficiency (P/O ratio, right) of liver mitochondria in the presence of 0, 10, or 25 μM ADP. Vehicle: ethanol. 3-Hydroxy stearic acid: 2 μM. $n = 3$. Data are shown as mean ± SD. Statistic: two-way ANOVA with Šídák's correct for multiple comparisons, $^{*}p < 0.05$; $^{**}p < 0.01$; $^{***}p < 0.001$

**B.** Absolute concentration of 3-hydroxy stearic acid in kidney (left), muscle (middle), and heart (right) from HFD-induced obese (DIO) mice supplemented with vehicle, stearic acid or 3-hydroxy stearic acid for two weeks. $n = 5$ per group. Data are shown as mean ± SD, individual values are presented. Statistic: unpaired *t*-test, $p < 0.05$; $^{**}p < 0.01$; $^{***}p < 0.001$.

**C-D.** Fat mass (**C**) and lean mass (**D**) of DIO mice supplemented with vehicle, stearic acid or 3-hydroxy stearic acid for two weeks. Body composition was assessed by EchoMRI. $n = 5$ per group. Data are shown as mean ± SD. Statistic: unpaired *t*-test.

**E.** Tissue weights of DIO mice supplemented with vehicle, stearic acid or 3-hydroxy stearic acid for two weeks. % of total body mass was presented. $n = 5$ per group. Data are shown as mean ± SD. Statistic: unpaired *t*-test.

**F.** Relative RNA expression level of ANT isoforms across 7 mouse tissues. Data was obtained from the BioGPS resource.

**G.** Dose titration of ANT2 inhibitor CATR on 3-OHSA-induced reduction in $H_2O_2$ production. $n = 3$ per group. Data are shown as mean ± SD. Statistic: unpaired *t*-test, $p < 0.05$; $^{**}p < 0.01$; $^{***}p < 0.001$.

**H.** Hepatic mitochondrial $H_2O_2$ production rate under basal condition and 1 μM CATR treatment. $n = 3$ per group. Data are shown as mean ± SD. Statistic: unpaired *t*-test.

**I-J.** Effects of the ANT inhibitors BA (20 μM) and CATR (1 μM), or the mPTP inhibitor CsA (1 μM), on the reduction of liver ΔΨm polarization rate induced by 2-hydroxystearic acid (**I**) or 3-OHFA12:0 (**J**). $n = 3$. Data are shown as mean ± SD, with individual values displayed. Statistic: unpaired t-test, $^{*}p < 0.05$; $^{**}p < 0.01$; $^{***}p < 0.001$.

**K.** mRNA expression of ANT2 in primary hepatocytes treated with siRNA targeting ANT2 or scrambled control. $n = 4$ per group. Data are shown as mean ± SD. Statistic: unpaired *t*-test, $p < 0.05$; $^{**}p < 0.01$; $^{***}p < 0.001$.

**L.** Effects of 3-OHSA on ΔΨm polarization rate in control and siRNA-ANT2 primary hepatocytes. $n = 3$ per group. Data are shown as mean ± SD. Statistic: unpaired *t*-test, $*p < 0.05$; $**p < 0.01$; $***p < 0.001$.
